# The Role of Visual Imagery in Face Recognition and Confidence Revisited: New Evidence from Aphantasia, Sampling Context Effects, and a Meta-Analysis

**DOI:** 10.64898/2026.07.01.734716

**Authors:** V. T. Königsmark, M-P. Stenner, R. R. Reeder, E. Azañón

## Abstract

The absence of voluntary visual imagery, known as aphantasia, offers a unique lens into the role of visual imagery in visual memory processes such as face recognition. While aphantasics often report difficulties, behavioral differences in standard tasks have generally been small. One possibility is that the contribution of visual imagery becomes apparent only when face recognition is especially demanding. We compared age- and gender-matched aphantasics and typical imagers on the more challenging long-form version of the *Cambridge Face Memory Test* (CFMT+) and on a measure of inverted face recognition. We also included tasks assessing object recognition and face perception. No group differences emerged for face perception or object recognition. By contrast, typical imagers outperformed aphantasics under high visual and mnemonic demands in face recognition in the laboratory cohort, particularly at the highest difficulty level of the CFMT+ and in inverted face recognition. This effect was attenuated or even absent in the online cohort. Drift-diffusion modelling indicated that this discrepancy was primarily driven by reduced response caution in online typical imagers. A meta-analysis of published short-form CFMT studies (total N = 432) revealed a moderate and reliable advantage for typical imagers (hedges *g* ≤ 0.41). Finally, across cohorts and tasks, aphantasics reported consistently lower subjective confidence, independent of accuracy. Overall, these findings suggest that visual imagery benefits face recognition, and highlight the need for caution in online testing, the predominant approach in aphantasia research.

## Introduction

The ability to recognize faces is essential for successful social interaction and relies on multiple sources of information, including visual and verbal processing (Bruce & Young, 1986; Haxby et al., 2000). Previous research has shown that visualizing a face from memory can improve subsequent recognition performance (Cabeza et al., 1997; Wu et al., 2012), with some studies proposing reactivation of perceptual representations in visual brain areas (Ishai et al., 2000). On the other hand, face recognition also benefits from semantic processing, i. e., through describing or labelling faces during encoding (Klatzky et al., 1982; McKelvie, 1976; Nakabayashi & Burton, 2008; Wickham & Swift, 2006).

The present study investigates whether the ability to generate visual imagery contributes to face recognition and confidence in individuals with aphantasia who experience a lack of voluntary visual imagery (Zeman et al., 2015). More broadly, we examine face recognition performance, confidence judgments, the robustness of observed effects across samples (online vs lab cohorts), and their overall strength across the literature. Aphantasia has been considered a natural knockout model (Dawes et al., 2022) for this type of investigation, as the absence of visual imagery allows for a direct test of its functional relevance in face processing. For individuals with typical imagery, it is tempting to assume that the ability to generate visual images should facilitate face recognition, as one might mentally compare a remembered stimulus with a currently perceived one (Keogh & Pearson, 2018). However, empirical evidence on whether individual differences in visual imagery provide a measurable advantage in face recognition remains mixed (Dance et al., 2023; McKelvie, 1994; Milton et al., 2021; Monzel et al., 2023b; Pounder et al., 2026).

McKelvie (1994) reported no advantage in face recognition with upright and inverted faces for individuals with more vivid imagery, when comparing “good” and “poor” visualizers. However, this study included external facial cues such as hair and accessories, possibly limiting its validity as a genuine test of internal-feature-based face recognition (see also Dance et al., 2023, for further discussion). The presence of these cues can lead to normal face recognition performance even in individuals with prosopagnosia (a condition marked by selective deficits in face recognition despite normal visual acuity and general cognitive functioning; Duchaine & Nakayama, 2004). In fact, Duchaine & Nakayama (2004) reported that 73% of 19 prosopagnosics performed in the normal range when such cues were present.

Milton et al. (2021) published the first empirical comparison of face recognition performance between individuals with aphantasia and typical imagers, finding no group differences in the *Famous Face Test*. However, this test uses highly familiar faces, which likely pose little challenge to perception and memory systems (Natu & O’Toole, 2011), likely due to the availability of richer and more discriminative representations compared to less familiar faces. In contrast, more recent work using the *Cambridge Face Memory Test* (CFMT; Duchaine & Nakayama, 2006), a well-established measure of unfamiliar face recognition, has reported group differences between aphantasics and typical imagers (Dance et al., 2023; Monzel et al., 2023b; Pounder et al., 2026). Monzel et al. (2023b) observed modest group differences when performance was averaged across the three CFMT levels, increasing in difficulty from recognizing faces shown in the same images as during encoding (same identities/same images), to recognizing faces after changing the lighting of images (same identities/novel images), or adding visual noise (same identities/novel and noisy images). They further proposed that these differences would become more pronounced as task difficulty increases, particularly when accuracy drops below approximately 52%. However, this prediction could not be directly tested within their dataset because, unlike the long form version of the CFMT, the standard CFMT does not include a near-chance level of difficulty.

The study addressed four related questions: (1) under what conditions visual imagery contributes to face recognition, (2) whether individuals with aphantasia show altered confidence or metacognitive patterns in task performance, (3) whether patterns of performance are robust to different sampling procedures, and (4) whether a meta-analysis of available literature on the CFMT supports a reliable association between visual imagery and face recognition ability.

To address the first aim, we examined whether the contribution of visual imagery to face recognition depends on task difficulty. Visual imagery is thought to support the generation and maintenance of detailed visual representations (Albers et al., 2013; Kosslyn et al., 2001; Pearson, 2019), which may aid the comparison of stored and currently perceived faces. When faces are easy to discriminate, recognition can often be based on salient visual cues (Duchaine & Weidenfeld, 2003). However, as perceptual ambiguity increases and faces become more difficult to distinguish, recognition is likely to depend more heavily on the quality of stored visual representations. If visual imagery contributes to these representations, then deficits associated with aphantasia should become increasingly apparent under more challenging recognition conditions. This prediction is consistent with evidence that performance differences tend to increase under higher task demands, such as high-precision visual working memory (Jacobs et al., 2018).

The extended CFMT+ (Russell et al., 2009) is well suited to test this prediction, as it progressively increases demands on both perceptual discrimination and memory retrieval, creating conditions under which visual imagery may play a compensatory role. The CFMT+ includes four levels of difficulty: the three standard CFMT levels plus an additional, more challenging, condition which was designed to identify “super-recognizers”, i.e., individuals with exceptionally high face recognition abilities (Russell et al., 2009). This final condition introduces five variations of the same target faces, including full profile views, emotional expressions, zoomed-in internal features and visible external features^1^.

In addition to overall task difficulty, we further examined whether visual imagery modulates featural and holistic processing in face perception. Face inversion is known to disrupt holistic face information (Valentine, 1988), resulting in coarser encoding of the spatial distances between individual features, and ultimately decreases recognition accuracy (Freire et al., 2000). Upright face recognition primarily engages holistic mechanisms (Richler et al., 2011). Nevertheless, greater reliance on individual feature processing has been associated with super-recognizers (Dunn et al., 2022), suggesting that enhanced processing of facial details can also facilitate recognition. Based on evidence that face imagery supports featural rather than holistic processing (Lobmaier & Mast, 2008), we hypothesized that typical imagers would outperform aphantasics when holistic cues are disrupted by inversion. This was tested by inverting faces of the CFMT-A (McKone et al., 2011), an Australian adaptation of the original test using new face identities compared to the upright CFMT+.

Our second aim examined group differences in subjective confidence and metacognitive evaluation. Prior work links visual imagery to confidence ratings (Reisberg & Leak, 1987; McKelvie, 1994; more recently: Liu & Bartolomeo, 2023; Monzel et al., 2023a; Wittmann & Şatırer, 2022), with more vivid imagers reporting higher confidence in imagery-related task performance than less vivid imagers, irrespective of actual performance differences (McKelvie, 1994; Reisberg & Leak, 1987). Similar results have been reported more recently in aphantasic samples (Wittmann & Şatırer, 2022) and differences in confidence have also been found in perceptual tasks that do not rely on direct assessments of imagery performance (Liu & Bartolomeo, 2023), possibly reflecting differences in self-perception in lower vividness groups (see Monzel et al., 2023a). Although aphantasics often report more face recognition difficulties (Zeman et al., 2020) with some reporting scores consistent with mild prosopagnosia (Dance et al., 2023), objective CFMT performance (commonly used to identify prosopagnosic traits) does not indicate increased prosopagnosia prevalence (Monzel et al., 2023b), suggesting a self-report-performance discrepancy. To examine this, we assessed trial-by-trial confidence ratings among aphantasics and typical imagers across both memory-based (CFMT+) and perceptual face tasks (*Cambridge Face Perception Test*: CFPT; Duchaine et al., 2007), as well as non-face control tasks (*Cambridge Bicycle Memory Test*: CBMT; and *Cambridge Car memory Test*: CCMT). We expected lower confidence in aphantasia to be independent of performance task type (similarly to weak imagery groups tested by McKelvie, 1994; Reisberg & Leak, 1987), potentially in alignment with reports that individuals with aphantasia often perceive themselves as cognitively “different” from others (Mawtus et al., 2024), a perception that may extend to tasks unrelated to visual imagery or task difficulty and which could be indicative of self-stigmatization.

Our third goal focused on whether group differences depend on sampling and testing environment. Given the low prevalence of aphantasia (1.5% - 4%; Beran et al., 2023; Dance et al., 2022), studies often rely on online recruitment (e.g., Dance et al., 2023; Liu & Bartolomeo, 2023; Monzel et al., 2023b), which might introduce variability in motivation and engagement. In particular, self-identified aphantasics may be motivated to participate by an interest in understanding their own cognitive experiences (see Mawtus et al., 2024). In contrast, typical imagers, who often serve as control participants, may be less intrinsically motivated and therefore less engaged with the task, potentially confounding group differences. To evaluate this, we compared lab and online data and examined their influence on accuracy, confidence, and decision-making behavior in the CFMT+ using drift diffusion modelling.

Finally, to address aim four, we conducted a meta-analysis of all currently available CFMT studies comparing aphantasics and typical imagers, including our data and datasets from Dance et al. (2023; https://osf.io/ny9bg), and Monzel et al. (2023; https://osf.io/k6bpf), both peer-reviewed, and Pounder et al. (2024; https://osf.io/f9jtr), which is openly available but not linked to a peer-reviewed article. All datasets employed only the three initial, less demanding levels of the CFMT+. This meta-analysis improves the precision of effect estimates across all currently available evidence, allowing us to determine whether group differences in face recognition emerge primarily under higher task difficulty and to clarify when visual imagery contributes.

In sum, the present study examined whether and under which conditions visual imagery contributes to face recognition, as well as how it relates to metacognitive evaluation in aphantasia. We expected a modest benefit of visual imagery under standard task conditions, increasing with task demands, alongside lower confidence ratings independent of performance. Finally, we examined whether these effects were robust across testing environments and integrated all available evidence in a meta-analysis to provide a more precise estimate of group differences. As reported below, our findings are consistent with a demand-dependent contribution of visual imagery to face recognition, alongside systematic reductions in reported confidence across both memory and perceptual tasks.

## Methods

### Participant cohorts

132 individuals (86 women, 46 men) aged 19 to 62 years (*M* = 34.1 years, *SD* = 10.5 years) completed the experiment either in the laboratory (lab cohort) or online (online cohort). Due to missing VVIQ data from one participant, the lab cohort comprised 28 aphantasics (21 women, 7 men; *M*_age_ = 31.8 years) and 27 age- and gender matched typical imagers (21 women, 6 men; *M*_age_ = 29.6 years), whereas 38 aphantasics (22 women, 16 men; *M*_age_ = 36.6 years) and 38 age- and gender matched typical imagers (22 women, 16 men; *M*_age_ = 36.6 years) participated in the online cohort. Aphantasics were recruited through the Aphantasia Research Database hosted at the Leibniz Institute for Neurobiology in Magdeburg, whereas typical imagers were recruited via word of mouth and snowball sampling (n = 13, 34.2%), two postings in Facebook groups for behavioral online study participation (n = 11, 29%), non-aphantasic contacts from our research database (n = 9, 23.7%) and the SONA Research Participation System at the University of Magdeburg (n = 5, 13.2%). Most participants were residents of Germany (n = 72), with additional individuals from Austria (n = 1), Switzerland (n = 1) and Spain (n = 2) and all were offered reimbursement (10€/h) via the Otto-von-Guericke University Magdeburg, unless voluntarily declined (5 aphantasics, 5 imagers). All participants gave their written or digital consent, and both studies were conducted in accordance with the Declaration of Helsinki (World Medical Association, 2013) and were approved by the ethics committee of the Otto-von-Guericke University Magdeburg (ethics number 118/22).

### Group classification

Aphantasic individuals and typical imagers were identified using a translated German (or Spanish) version of the *Vividness of Visual Imagery Questionnaire* (VVIQ; Marks, 1973), which asks participants to rate the vividness of visual images evoked by 16 items spanning four different scenes (e.g., “*The sun is rising above the horizon into a hazy sky”*, Item 5). Response options range from 1 (*“No image at all, you only “know” that you are thinking of the object”*) to 5 (*“Perfectly clear and as vivid as normal vision”*), yielding total scores between 16 and 80. Lab-based aphantasics scored between 16 and 26 on the VVIQ (*M*_VVIQ_ = 18.8, *SD* = 3.27), and online aphantasics between 16 and 21 (*M*_VVIQ_ = 16.6, *SD* = 1.39). Consistent with previous classifications (Dance et al.2023; Monzel et al., 2023b), they reported to have no image at all, or at most described their experience as dim or vague. Typical imagers scored between 32 and 73 (*M*_VVIQ_ = 57.8, *SD* = 10.6), and between 32 and 80 (*M*_VVIQ_ = 59.8, *SD* = 14) in the lab- and online cohort respectively. Among the latter, five individuals scored above 75, a range previously associated with “hyperphantasia” (Milton et al., 2021; Reeder et al., 2024; Zeman et al., 2020).

In line with prior research (Bainbridge et al., 2021; Weber et al., 2024), a subset of our participants from both cohorts (n = 112, 57 aphantasics, 55 typical imagers) also completed the *Object-Spatial-Imagery and Verbal Questionnaire* (OSIVQ; Blazhenkova & Kozhevnikov, 2008), assessing individual differences in object imagery, spatial imagery and verbal cognitive styles. We observed significant differences in object imagery between self-reported aphantasics and typical imagers (*t*(110) = 17.8, *p* < .0001, *d* = 3.4), consistent with VVIQ results (see Figure 1C), but found no notable differences in spatial imagery or verbal thinking (both t ≤ 1.36; both p > .18).

**Figure 1.**
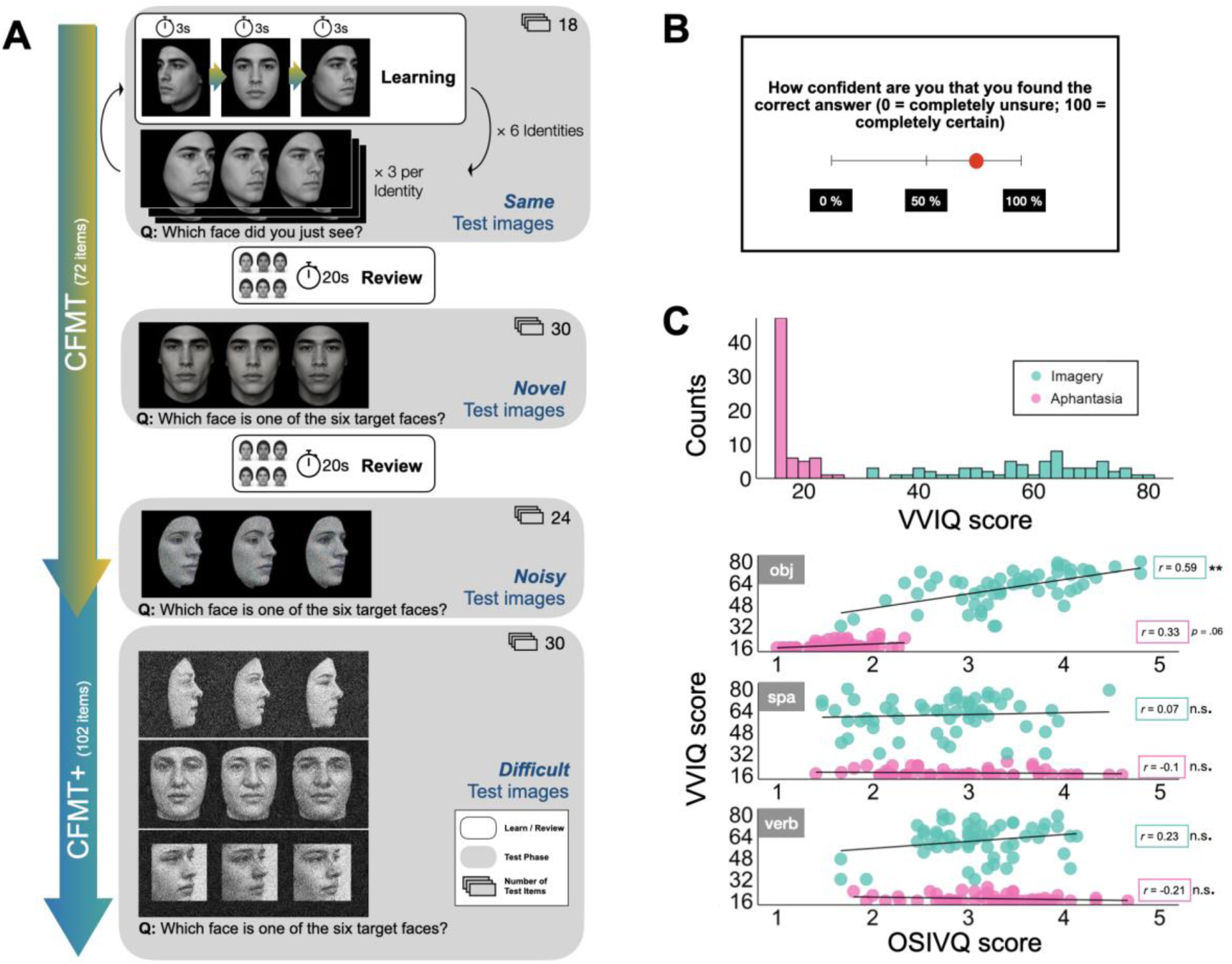
Task design: Individual levels, response items and task questions of the CFMT and CFMT+ (A) and trial-by-trial response cues for confidence ratings (B). Group counts of VVIQ scores and correlations with mean object, spatial and verbal OSIVQ scores (C), *r* = Pearson’s correlation coefficient, all *p*-values Holm corrected. Face images shown are AI-generated and are included for illustrative purposes only. They were not used as experimental stimuli and do not depict real individuals.

### Experimental tasks

The procedures and trial order of all tasks follow the original protocols reported in the respective task publications and were implemented as in previous studies (Duchaine & Nakayama, 2006; Dance et al., 2023; Monzel et al., 2023b). Crucially, we extended the original protocols by obtaining confidence ratings after every trial, allowing us to assess participants’ metacognitive judgments alongside objective performance. Precise information about display configurations for each task as well as size of stimuli can be found in the supplemental material.

#### CFMT+

All participants completed the extended, or “long form,” version of the *Cambridge Face Memory Test* (CFMT+; Russell et al., 2009), developed to assess super-recognizers (individuals with exceptionally high face recognition ability). The CFMT+ comprises four levels of increasing difficulty (henceforth “levels”: same images, novel images, noisy images, and difficult images (Figure 1A). The first three levels correspond to the original (“short form”) CFMT (Duchaine & Nakayama, 2006), whereas the fourth level contains additional highly challenging recognition trials.

During the initial learning phase, participants were introduced to six unfamiliar Caucasian male identities. Each identity was presented sequentially from three viewpoints (frontal, left three-quarter, and right three-quarter view), with each viewpoint displayed for 3 s and separated by a 0.5 s interstimulus interval. Presenting multiple viewpoints encouraged learning of the facial identity rather than memorization of a specific image. Immediately after learning an identity, participants completed three three-alternative forced-choice (3AFC) recognition trials. On each trial, three faces were presented simultaneously, and participants identified the face they had just learned by pressing one of three number keys (1, 2, or 3) on the keyboard. One face was the target identity and the other two were unfamiliar distractor identities. Within each trial, all faces were shown from the same viewpoint, and across the three trials the target identity was tested once from each of the three studied viewpoints. Importantly, the target face was presented using the identical image shown during learning. Across the six identities, this resulted in 18 recognition trials, constituting the *same-images* level.

Following the *same-images* level, participants completed a review phase in which all six target identities were presented simultaneously in frontal view for 20 s, arranged in a 2 × 3 grid. Participants then completed the *novel-images* level (30 trials), followed by a second 20 s review phase and the *noisy-images* level (24 trials). Finally, participants completed the *difficult-images* level (30 trials) without an additional review phase.

The three latter levels followed the same procedure. Each trial consisted of a three-alternative forced-choice (3AFC) recognition task in which participants identified which of three faces corresponded to one of the six identities learned during the initial study phase. In the novel-images phase, test faces were shown from previously unseen viewpoints and/or under novel lighting conditions. In the noisy-images phase, images were further degraded using Gaussian visual noise. Finally, the difficult-images phase (30 trials) introduced five challenging variations of each target face, including full-profile views, smiling expressions, zoomed-in internal features, visible external features, and additional emotional expressions, all presented under increased visual noise. Whereas the short form CFMT uses many novel distractor identities across trials, the long form makes the task harder by reusing a smaller set of distractors more frequently, thereby increasing distractor familiarity and reducing reliance on simple target familiarity (for further methodological details, see Russell et al., 2009). In the present study, a trial-by-trial confidence rating procedure was introduced as an extension to the original CFMT and CFMT+ protocols. This procedure was also applied to the remaining tasks (see below). Specifically, after each test trial, participants rated the confidence that their decision was correct using a continuous visual analog slider ranging from 0% (“completely uncertain”) to 100% (“completely certain”; note that the original was in German or Spanish; Figure 1B). Confidence ratings were provided using the mouse. Both the 3AFC responses and confidence judgments were non-speeded. The face array remained visible until a response was made. Across levels participants performed 102 trials.

#### Modified CFMT-A

To assess recognition of inverted faces, all participants completed a modified version of the Australian CFMT (CFMT-A; McKone et al., 2011). We chose the CFMT-A as it employs the same structure as the CFMT but uses a different stimulus set to prevent memory carryover from upright to inverted trials. We modified the original version by rotating upright images 180° (Modified CFMT-A, henceforth “CFMT-AM”), hence presenting all images upside-down. Unlike the CFMT+, however, the CFMT-A comprises only the three standard CFMT levels (*same images*, *novel images*, and *noisy images*; n total trials = 72). Trial-by-trial confidence ratings were also added here.

#### CBMT/CCMT

To measure general object recognition from memory, participants additionally completed either the design-matched *Cambridge Bicycle Memory Test* (CBMT; Dalrymple et al., 2017) or the design-matched *Cambridge Car Memory Test* (CCMT; Dennett et al., 2012). The CBMT was used in the lab setting, but due to observed ceiling effects (i.e., accuracy rates exceeding 85% across all difficulty levels and groups), the CCMT was used for online participants. Both the CBMT and CCMT follow the same basic structure of the CFMT described above, showing either bicycles or cars and yielding a total of 72 trials. As in the CFMT+, trial-by-trial confidence ratings were also added here.

#### CFPT

Lastly, all participants completed the Cambridge Face Perception Test (CFPT; Duchaine et al., 2007), a face-matching task designed to assess perceptual processing of facial identity while minimizing memory demands. On each trial, a target face was presented at the top of the screen in a three-quarter view. Below the target, six frontal-view faces were displayed simultaneously in a horizontal row. These six faces were morphs between the target identity and another identity and therefore varied systematically in their similarity to the target face (e.g., 88%, 76%, 64%, 52%, 40%, and 28% target identity). Participants were instructed to arrange the six morphs from most similar to least similar to the target by dragging and dropping them into the desired order. The target face remained visible throughout the trial, allowing participants to make direct perceptual comparisons rather than relying on memory. Once satisfied with their ranking, participants submitted their response (self-paced; maximum response window = 60 s). The task comprised 16 trials derived from eight target identities (plus two practice trials). Each identity appeared once in an upright orientation and once in an inverted orientation, with upright and inverted trials interleaved throughout the task. Performance was scored by summing the absolute deviations of each face’s assigned position from the correct morph order. For example, placing the face with the highest similarity to the target (88% overlap) in the third position yielded a deviation score of 2. To assess metacognition, we added the same trial-by-trial confidence measure as before (Figure 1B).

### Questionnaires

All participants completed German-translated (or Spanish) versions of five questionnaires, including the VVIQ, the PSI-Q (Andrade et al., 2014), the OSIVQ (Blazhenkova & Kozhevnikov, 2009), the SAM (Palombo et al., 2013), and a custom post-hoc questionnaire regarding participant’s subjective experiences and used strategies in each test.

The VVIQ (Marks, 1973) is a widely used self-report measure of imagery vividness, in which participants rate the clarity of mental images across everyday scenarios. Ratings are typically made on a 5-point scale ranging from 1 (“no image at all”) to 5 (“perfectly clear and vivid”), and the measure is commonly used to characterize individual differences in visual imagery, including aphantasia. The PSI-Q measures the vividness of sensory imagery across seven modalities: vision, sound, smell, taste, touch, bodily sensations, and emotions, e.g., *“Imagine feeling in love”*, on an 11-point Likert scale (0 = not vivid at all, 10 = extremely vivid). To assess individual differences in cognitive styles, including object imagery, spatial imagery, and verbal thinking, the OSIVQ was administered. This questionnaire consists of statements such as *“I enjoy pictures with bright colors and unusual shapes like the ones in modern art”* (object scale), “*I was very good in 3D geometry as a student”* (spatial scale), or *“I have difficulty expressing myself in writing”* (verbal scale), which participants rate on a 5-point scale (1 = “totally disagree”, 3 = “not sure”, 5 = “absolutely agree”). Lastly, we administered the SAM to assess participants’ degree of subjectively retained autobiographical memories. The SAM evaluates episodic, semantic, spatial, and future-oriented dimensions of autobiographical memory on a 5-point Likert scale ranging from 1 (“strongly disagree”) to 5 (“agree strongly”; see Supplementary Figure 4, section 2.4.1). Lastly, participants responded to a post-questionnaire assessing additional demographics, daily-life imagery experiences and strategy-use throughout the experiment, i.e., “*Did you try to remember the entire image or focus on specific parts? Please describe your strategy in more detail.*” (see Supplementary Figure 5 and Figure 6, section 2.4.2).

### Procedure

All participants completed the VVIQ and PSI-Q at least 24 hours before their experimental session. Prior to the experiment, participants received standardized task instructions (in-person for lab participants; via video tutorial for online participants). Participants then completed four tests in a fixed order: (1) upright face recognition (CFMT+), (2) object recognition (CBMT or CCMT), (3) inverted face recognition (CFMT-AM) and (4) face perception (CFPT) with untimed breaks between tasks. The full session lasted approximately 75 minutes.

Upon completion, participants filled in the post-experiment questionnaire assessing additional demographics, subjective experiences and strategy use (see Supplementary Figure 5 and Figure 6). The OSIVQ (Figure 1) and SAM (see Supplementary Figure 4, section 2.4.1) were administered after the experimental session. The experiment was not publicly accessible; study links and instructions were distributed individually following multiple communications with the experimenter to verify eligibility and retain procedural control.

### Apparatus

The experimental battery was implemented in MATLAB (2021b) using Psychtoolbox-3 (MathWorks Inc., Natick, MA, USA) for the lab cohort and adapted to PsychoPy (Version 2024.1.1; Peirce et al., 2024) for the online cohort.

Lab participants completed the study on a desktop computer in a controlled testing environment. Stimuli were presented on an ASUS VG248QE 24-inch monitor (1920 × 1080 pixels, 60 Hz), at an approximate viewing distance of 50 cm. Stimulus sizes and layouts followed the validated formats of the original tasks.

Online participants were instructed to complete the experiment under similar conditions (quiet environment, no multitasking, adequate lighting). To approximate lab-based stimulus presentation, a credit-card calibration procedure (Morys-Carter, 2023) was used to scale stimuli to differences in physical screen size. Participants were instructed to maintain viewing distance of approximately 50 cm, although this could not be verified. Due to ceiling effects of the CBMT in the lab cohort, the CCMT was administered instead in the online cohort.

### Statistical analyses

All analyses were conducted in JASP (v0.19.3; JASP Team, 2025), RStudio (R 4.4.2 software; R Core Team, 2024) and Python (v3.6; Python Software Foundation, 2016; for drift diffusion modelling). Primary outcome measures were recognition accuracy (percentages correct) and response times (RTs; seconds from stimulus onset to key press). Given the positive skew typical of reaction time (RT) distributions, even in non-speeded tasks (Luce, 1991; Van Zandt, 2000), median RTs were analyzed (see Ratcliff, 1993; Whelan, 2008).

Recognition performance for upright faces was assessed using mixed ANOVAs with Group (typical imagers, aphantasics) as a between-subject factor and Item Difficulty (same-images, novel-images, noisy-images; adding difficult-images for the CFMT+) as a within-subject factor. Where appropriate, we integrate both cohorts by introducing *Cohort* (Lab vs. Online) as an additional between-subjects factor. For repeated-measures factors, violations of sphericity were addressed using the Huynh–Feldt correction, and corrected degrees of freedom are reported where applicable.

CFPT performance was indexed by the sum of absolute deviations from the correct face order, with lower scores indicating better performance (see above). These scores were analyzed using a 2 × 2 × 2 ANOVA with Group, Cohort, and Orientation (upright, inverted) as factors.

To examine potential speed-accuracy trade-offs, accuracy and RT were jointly modelled using a drift diffusion model (see next paragraph).

In addition to frequentist analyses, results were evaluated within a Bayesian framework using Bayes Factors (BFs), interpreted according to established conventions (Quintana & Williams, 2018), with BF_10_ classified as anecdotal (1–3), moderate (3–10), strong (10–30), very strong (30–100), or extreme (>100). BF_10_ < 1 suggest evidence in favor of the null hypothesis. To ensure reproducibility, a fixed random seed of 42 was used for all sampling-based procedures. Robustness was assessed across a range of prior widths; results were consistent across specifications (see Supplementary Material, Section 3.3).

#### Hierarchical drift diffusion model

To examine group differences in latent decision processes beyond accuracy and response times, we applied the drift diffusion model (DDM; Ratcliff, 1978). The DDM conceptualizes decisions as noisy evidence accumulation over time toward a response boundary and is parameterized by (1) drift rate (*v*), reflecting the average speed and direction of information accumulation, (2) boundary separation (*a*), indicating the amount of information required before committing to a response (i.e., response caution), (3) non-decision time (*t*_0_), capturing perceptual and motor delays, and (4) the starting point (*z*), representing *a priori* bias toward one response option.

The DDM is traditionally applied to binary decision tasks with two physical stimuli (e.g., “old” vs. “new”) and was successfully used in non-speeded tasks (Myers et al., 2022) and in face recognition paradigms, specifically (e.g., Williams et al., 2023; Powell et al., 2019). In our case, each three-alternative forced choice (3AFC) trial can conceptually be reduced to a binary classification at the level of accuracy: a response is either “old” (target / correct) or “new” (distractor / wrong), regardless of which “new” item was selected. While this simplification does not map directly onto the classic 2AFC DDM, it provides a viable framework for estimating underlying decision parameters (see Voss et al., 2015 for a discussion on adapting DDM to non-binary tasks). Thus, we treated the decision as a binary process (target vs. distractor) for modeling purposes. The resulting parameter estimates are therefore not directly comparable to those derived from true binary paradigms; however, they remain valid for within-study (i.e., cohort) comparisons, allowing us to investigate how decision dynamics vary between groups and across conditions.

DDM parameter estimation requires an error rate above 2.5% (Heathcote et al., 2018), to maintain model stability and avoid bias from uninformative accuracy distributions. We therefore excluded the same-images condition in the CFMT+ from the analysis because its error rate fell below 2.5%. Given the limited number of trials across difficulty levels that ranged between 18 and 30, we estimated DDM parameters using a hierarchical Bayesian framework via the Hierarchical Drift Diffusion Model (HDDM) toolbox (Wiecki et al., 2013) in Python 3.6 (Python Software Foundation, 2016). This approach is particularly advantageous in tasks with relatively few trials per condition (Ratcliff & Childers, 2015). Being hierarchical in nature, the HDDM simultaneously estimates parameters at the group (either aphantasics or typical imagers) and individual level, borrowing statistical strength across participants. Posterior distributions are sampled using Markov Chain Monte Carlo (MCMC) procedures based on the Wiener first-passage time distribution (Wald, 1947; Feller, 1968).

We ran separate HDDM models for the upright and inverted face recognition tasks, allowing the drift rate (*v*) and boundary separation (*a*) to vary by group and individual. The non-decision time (*t₀*) and starting point (*z*) parameters were included in the models but were held constant across groups, following standard modeling practice unless bias or motor effects are hypothesized (Wiecki et al., 2013). Drift-rate and boundary separation parameters were set as free unless otherwise specified and estimated using default non-informative priors provided by the HDDM toolbox. In line with HDDM defaults, 5% of trials were assumed to reflect outlier responses (e.g., attentional lapses or accidental responses) and were modeled as arising from a separate random-response process rather than the diffusion process itself. We initially ran models with 10,000 posterior samples and 200 burn-in iterations. However, a comparison with a shorter chain using 2000 samples and 20 burn-ins showed no meaningful difference in posterior estimates, suggesting that the posterior distributions were stable and adequately sampled. The shorter chain was used in the final reported analysis for computational efficiency. To reduce autocorrelation among consecutive samples in the posterior distribution, we kept every fifth sample (thinning factor = 5), effectively discarding the immediate ones to ensure the retained samples were more independent. Model convergence was assessed using the Gelman-Rubin R-hat statistic (Gelman & Rubin, 1992), which produced chain values < 1.05, suggesting good convergence.

#### Meta-analysis

We conducted three random-effects meta-analyses (Borenstein et al., 2009), one per difficulty level, using the *metafor* package (Viechtbauer, 2010) in RStudio (R 4.4.2 software).

Given observed cohort differences, samples were treated separately by testing context, including the present lab and online cohorts, fully online samples from Monzel et al. (2023b) and Pounder et al. (2024; dataset), and both lab-based and remote cohorts (via Skype calls) from Dance et al. (2023)^2^. A random-effects model was used to account for between-sample variability in participant populations and testing conditions. Effect sizes were computed as Hedges’ *g* (Borenstein et al., 2009).

## Results

Results are reported separately for the laboratory and online cohorts, which were collected as independent studies (laboratory: 2021 and 2023; online: 2024). Cohort (lab vs. online) was included as an additional between-subjects factor in analyses that combined participants from both samples. This approach reflects observed cohort-related differences in performance, particularly between the lab and online control group.

### Face recognition from memory with upright faces (CFMT+, CFMT)

#### Study 1: Lab cohort

A 2 × 4 mixed-design ANOVA on accuracy (percentages correct) with Group (typical imagers, aphantasics) and Item Difficulty (same, novel, noisy, difficult images) revealed a main effect of Item Difficulty, *F*(2.67, 141.7) = 230.3, *p* < .001, *η^2^*_p_ = 0.81, BF_10_ > 1000, indicating decreasing accuracy with increasing difficulty. There was also a main effect of Group, *F*(1, 53) = 8.09, *p* = .006, *η^2^*_p_ = 0.13, BF_10_ = 15.2, with aphantasics performing worse (*M* = 66.8%, *SD* = 12.2%) than typical imagers (*M* = 75.9%, *SD* = 10.7%).

The Group × Item Difficulty interaction was significant, *F*(2.77, 146.8) = 3.42, *p* = .023, *η^2^*_p_ = 0.06, BF_10_ = 7.5. Holm-corrected post hoc comparisons showed no reliable group differences for *same* (level 1) or *noisy* (level 3) images, (both *p* = .096, both *d* ≤ 0.46, both BF_10_ ≤ 1.65), moderate evidence for a group difference in the *novel*-images condition (level 2), *t*(53) = 2.26*, p* = .04, *d* = 0.61, BF_10_ = 4.21, and strong evidence in the *difficult*-images condition (level 4), *t*(53) = 3.73*, p* < .001, *d* = 1, BF_10_ = 116 (see Figure 2A).

**Figure 2.**
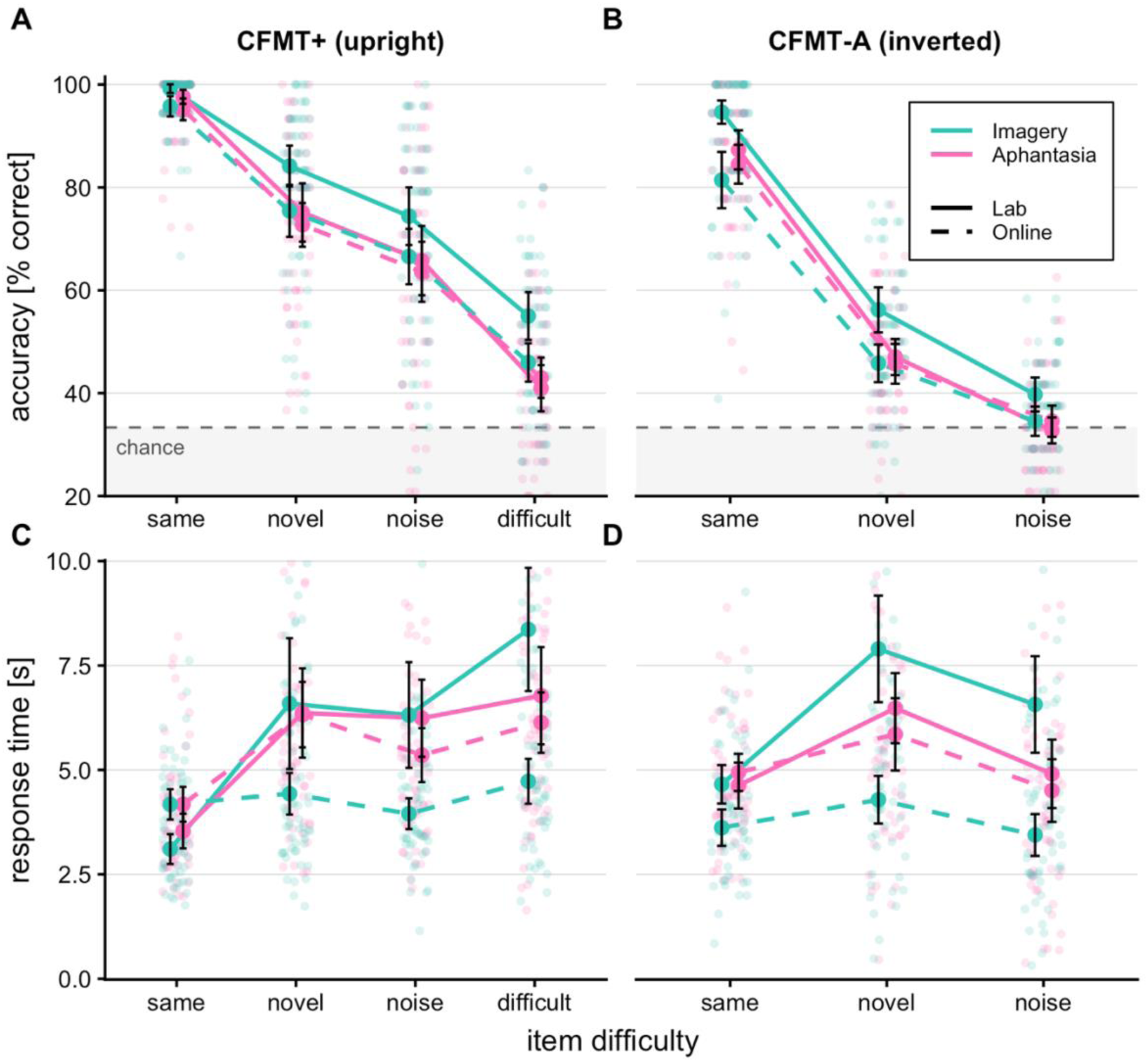
Accuracy and response times (RTs). Top row: face recognition accuracy for upright (CFMT+) (A) and inverted (CFMT-A) (B) faces, shown as mean percentage of correct responses per test condition. Bottom row: Group mean response times (in seconds), calculated from individual median response times, for upright (C) and inverted (D) test conditions. Colors (group) and line types (cohort): turquoise = Typical Imagers, magenta = Aphantasics; solid line = lab participants, dashed line = online participants. Error bars signify standard errors of the mean (SEM). For both accuracy and response time measures, aphantasic participants (depicted in pink) showed consistent performance across cohorts (laboratory vs. online), whereas imagery participants (depicted in blue) exhibited cohort-dependent differences in performance.

For comparability with existing CFMT literature, we additionally analyzed the short form (three levels; see Supplementary Table 5, section 1.1).

### Study 2: Online cohort

The same analysis was conducted online (38 typical imagers, 38 aphantasics). A main effect of Item Difficulty was observed, *F*(2.54, 187.7) = 306.28, *p* < .001, *η^2^*_p_ = 0.81, BF_10_ > 1000, but unlike study 1 conducted in the lab, there was no main effect of Group, (*p* = .397, BF_10_ = 0.25), nor interaction (*p* = .84, BF_10_ = 0.05).

### Group vs. cohort analyses: Accuracy

To formally assess differences between the laboratory and online cohorts, we conducted a 2 × 2 × 4 ANOVA (Cohort × Group × Item Difficulty). This confirmed a main effect of Item Difficulty (*F*(2.54, 323) = 521.39, *p* < .001, *η^2^*_p_ = 0.80, BF_10_ > 1000). Evidence for the remaining effects differed across inferential frameworks. Although the frequentist analyses yielded significant effects (Group × Item Difficulty, *p* = .048; Cohort, *p* = .038; Group, *p* = .010), the corresponding Bayes factors provided only anecdotal or inconclusive evidence (BF_10_ = 0.58, 0.73, and 1.67, respectively). Post-hoc tests showed a group difference only at the highest difficulty (*p_uncorr_* = .005, *p_corr_* = .02, *d* = 0.54, BF_10_ = 27.56), (all other levels: *t*(129) ≤ 1.91, all *p_corr_*≥ .09; see Table 2; all accuracy means and SDs are reported in Supplementary Table 1, section 1.1).

Critically, there was no Cohort × Group interaction, *F*(1, 127) = 2.20, *p* = .141, BF_10_ = 0.64. Exploratory comparisons showed reduced accuracy for typical imagers in the online relative to the lab cohort (*M*_Lab_ = 75.9%, *M*_Online_ = 68.3%, *t*(63) = −2.51, *p* = .015, BF_10_ = 3.47), whereas aphantasics performed comparably across cohorts (*M*_Lab_ = 66.8%, *M*_Online_ = 65.8%, *t*(64) = 0.36, *p* = .72, BF_01_ = 3.72). Although this pattern should be interpreted cautiously given the absence of a significant interaction, it is consistent with reduced engagement in the online typical-imagery group, which we examine further using RT and drift-diffusion analyses below.

### Group vs. cohort analyses: Response times

To further examine cohort differences, a 2 × 2 × 4 ANOVA on median RTs (Cohort × Group × Item Difficulty) was conducted. RTs increased with Item Difficulty, *F*(2.4, 305.4) = 73.64, p < .001, BF10 > 1000, but this effect was qualified by a significant three-way interaction, *F*(2.4, 305.4) = 6.62, *p* = .001, BF_10_ = 125.5. Given this interaction, lower-order effects are not interpreted further.

To probe the three-way interaction, we analyzed individual RT linear slopes across difficulty levels. A 2 × 2 ANOVA (Cohort × Group) on slopes revealed a main effect of Cohort, *F*(1, 127) = 38.27, *p* < .001, *η^2^*_p_ = 0.23, BF_10_ > 1000, qualified by a Cohort × Group interaction, *F*(1, 127) = 9.18, *p* = .003, *η^2^*_p_ = 0.07, BF_10_ = 8.6. Follow-up tests showed increasing RTs with difficulty in all groups (all p < .001, all BF_10_s > 74), except for the online typical-imagery group, which showed no reliable modulation, (*p* = .07, BF_10_ = 0.82; all RT means and SDs in Supplementary Table 2, section 1.1). The pattern of relatively flat RT slopes combined with reduced accuracy suggests diminished sensitivity to task demands in the online control group (Figure 2C). We return to this issue in the decision-model analyses below.

### Group vs. cohort analyses: Hierarchical drift diffusion modeling

#### Response caution: boundary separation

Given the concerns about reduced task engagement in the online imagery group, we first examined boundary separation as an index of response caution using hierarchical drift diffusion modeling (excluding the same-images condition due to low error rates; Heathcote et al., 2018; 2.38% for aphantasics, 0.79% for typical imagers). A 2 × 2 × 3 ANOVA revealed main effects of Item Difficulty and Cohort (both *p* < .001, BF_10_ > 1000) but no clear main effect of Group (*p* = .18, but moderate evidence, BF_10_ = 3.42). Critically, these factors were qualified by a Cohort × Group interaction, *F*(1, 127) = 7.29, *p* = .008, *η^2^*_p_ = 0.05, BF_10_ = 7.48. Follow-up comparisons showed that typical imagers tested online (*M* = 5.15, *SD* = 1.04) exhibited reduced response caution relative to the lab cohort (*M* = 5.15, *SD* = 1.04), *t*(63) = −5.37 *p* < .001, *d* = 1.36, BF_10_ > 1000), whereas aphantasics did not differ across cohorts (*p* = .24, BF_10_ = 0.46). This pattern is consistent with reduced task engagement or a shift toward a less cautious response strategy in the online imagery group.

### Evidence accumulation: drift rates

Drift rate indexes the efficiency of evidence accumulation. A corresponding ANOVA revealed a strong main effect of Item Difficulty, *F*(1.91, 243.6) = 415.99, *p* < .001, *η^2^*_p_ = 0.77, BF_10_ > 1000, confirming that evidence accumulation declined with increasing task difficulty (Myers et al., 2022), and a main effect of Group, *F*(1, 127) = 14.41, *p* < .001, *η^2^*_p_ = 0.10, BF_10_ = 55.62. No effects involving Cohort were observed (all *p* ≥ .13, BF_10_ ≤ 0.51). Thus, despite reduced accuracy and lower response caution in online typical imagers, drift rates were preserved, indicating intact evidence accumulation. This pattern suggests a shift toward faster, less cautious responding rather than a deficit in perceptual or decisional processing. Follow-up comparisons confirmed that typical imagers showed higher drift rates than aphantasics across all analyzed CFMT+ difficulty levels (all *p_corr_* ≤ .007, BF_10_ ≥ 5.24).

### Face recognition from memory with inverted faces (CFMT-A)

To assess recognition under disrupted holistic processing, participants completed an inverted-face task using novel stimuli from a modified Cambridge Face Memory Test (CFMT-AM). Faces were presented upside down to impair holistic processing, and participants selected, again, the target from three alternatives (chance = 33.33%).

#### Study 1: Lab cohort

In the lab cohort, a 2 × 3 ANOVA (Group × Item Difficulty: same, novel, noisy) on accuracy revealed a main effect of Item Difficulty, *F*(1.92, 101.9) = 532.31, *p* < 0.001, *η^2^*_p_ = 0.91, BF_10_ > 1000, and Group, *F*(1, 53) = 19.29, *p* < .001, *η^2^*_p_ = 0.27, BF_10_ = 191.2, with typical imagers (*M* = 60.3%, *SD* = 8.16%) outperforming aphantasics (*M* = 52.3%, *SD* = 5.77%). There was no Group × Item Difficulty interaction, (*p* = .80, BF_10_ = 0.48). Holm-corrected post hoc comparisons confirmed that typical imagers outperformed aphantasics across all levels (all *p* ≤ .006, all BF_10_ ≥ 13.51; see Figure 2B).

#### Study 2: Online cohort

In contrast to the lab cohort, the online cohort showed no accuracy differences between groups for inverted faces, mirroring the pattern observed for upright faces. The ANOVA revealed a strong main effect of Item Difficulty, *F*(1.79, 132.1) = 323.67, *p* < .001, *η^2^*_p_ = 0.81, BF_10_ > 1000, but no main effect of Group, (*p* = .62, BF_10_ = 0.16), and no interaction (*p* = .64, BF_10_ = 0.08).

#### Group vs. cohort analyses: Accuracy

To examine cohort-related differences, a 2 × 2 × 3 ANOVA (Cohort × Group × Item Difficulty) was conducted on inverted-face accuracy (all accuracy means and SDs in Supplementary Table 1, section 1.1). A strong effect of Item Difficulty was observed, *F*(1.91, 242.9) = 750.8, *p* < .001, *η^2^*_p_ = 0.86, BF_10_ > 1000, along with main effects of Group, *F*(1, 127) = 6.01, *p* = .016, *η^2^*_p_ = 0.05, BF_10_ = 3.68, with typical imagers showing higher accuracy than aphantasics, and Cohort, *F*(1, 127) = 13.8, *p* < .001, *η^2^*_p_ = 0.10, BF_10_ = 131.3, with higher accuracy in the lab than online. Critically, a Cohort × Group interaction emerged, *F*(1, 127) = 10, *p* = .002, *η^2^*_p_ = 0.07, BF_10_ = 11.32, indicating that group differences varied across cohorts. The Cohort × Item Difficulty interaction was not significant, (*p* = .058, BF_10_ = 1.56). Based on our earlier findings on upright face recognition, we next examined RTs.

#### Group vs. cohort analyses: Response times

RTs showed a main effect of Cohort, *F*(1, 127) = 14.41, *p* < .001, *η^2^*_p_ = 0.1, BF_10_ > 1000, qualified by a Group × Cohort interaction, *F*(1, 127) = 9.87, *p* = .002, *η^2^*_p_ = 0.07, BF_10_ = 16.42. There were also a Group × Item Difficulty interaction, *F*(1.83, 232.7) = 3.38, *p* = .04, *η^2^*_p_ = 0.03, BF_10_ = 1.69, and a Cohort × Item Difficulty interaction, *F*(1.83, 232.7) = 12.97, *p* < .001, *η^2^*_p_ = 0.09, BF_10_ > 1000. The three-way interaction did not reach significance, (*p* = .07), although Bayesian evidence suggested a moderate effect (BF_10_ = 4.25).

Follow-up comparisons mirrored the pattern observed for upright faces: aphantasics showed no cohort differences (*p* = .64, BF_10_ = 0.28), whereas typical imagers responded faster online (*M* = 3.78s, *SD* = 1.5s) than in the lab (*M* = 6.3s, *SD* = 2.62s; *t*(64) = −4.91, *p* < .001, *d* = 1.23, BF_10_ > 1000; Figure 2D; all RT means and SDs in Supplementary Table 2, section 1.1).

#### Group vs. cohort analyses: Hierarchical drift diffusion modeling

Consistent with RT results, boundary separation showed a main effect of Cohort, *F*(1, 127) = 18.48, *p* < .001, *η^2^*_p_ = 0.13, BF_10_ > 1000, but no main effect of Group (p = .91; although with moderate evidence for an effect, BF_10_ = 6.09). Importantly, this effect was modulated by a Cohort × Group interaction, *F*(1, 127) = 11.37, *p* = .001, *η^2^*_p_ = 0.08, BF_10_ = 22.98. Follow-up comparisons indicated reduced response caution in the online imagery group relative to the lab group, *t*(63) = 5.37, *p* < .001, *d* = 1.35, BF_10_ > 1000, whereas, again, no differences were observed for aphantasics, (*p* = .51, BF_10_ = 0.31; Figure 3D).

**Figure 3.**
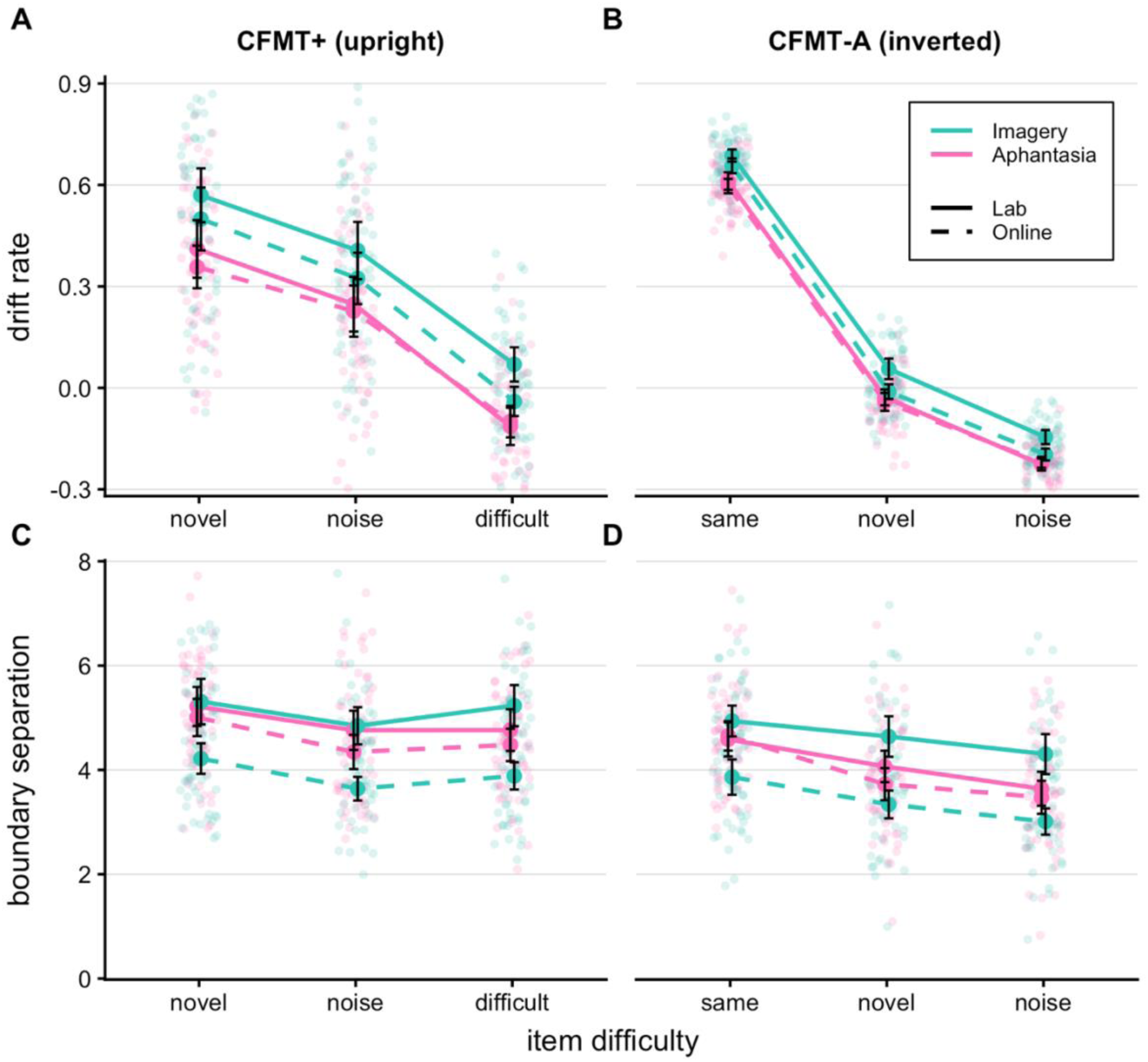
Drift diffusion parameters. Evidence accumulation shown as mean estimated drift rate values (*v*) (top row: (A) = upright, (B) = inverted), and response caution, shown as mean estimated boundary separation values (*a*) (bottom row: (C) = upright, (D) = inverted) per test condition of upright (CFMT+) and inverted (CFMT-AM) face recognition. Note that the same-images condition was excluded from parameter estimation for upright conditions due to an error rate < 2.5%. Colors (group) and line types (cohort): Turquoise = Typical Imagers, magenta = Aphantasics; solid line = lab participants, dashed line = online participants. Error bars signify standard errors of the mean (SE).

Drift rate (evidence accumulation; Figure 3B) was analyzed using a 2 x 2 x 3 ANOVA (Cohort × Group × Item Difficulty). Drift rates decreased with Item Difficulty, *F*(2, 257.6) = 5929, *p* < .001, *η^2^*_p_ = 0.98, > 1000, and differed by Group, *F*(1, 127) = 48.35, *p* < .001, *η^2^*_p_ = 0.28, BF_10_ > 1000. In contrast to the upright task, there was also a main effect of Cohort, *F*(1, 127) = 11.14, *p* = .001, *η^2^*_p_ = 0.08, BF_10_ = 21.36 and a Cohort × Group interaction, *F*(1, 127) = 6.09, *p* = .0015, *η^2^*_p_ = 0.05, BF_10_ = 4.88. Follow-ups showed that this interaction was driven by lower drift rates in the online imagery group relative to the lab imagery group, *t*(63) = 3.92, *p* < .001, *d* = 0.99, BF_10_ = 110.3, with no corresponding cohort differences in aphantasics (*p* = .52, BF_10_ = 0.3). This suggests less efficient evidence accumulation in the online imagery group in the inverted task, which is overall more demanding than the upright condition. More generally, this pattern may reflect reduced task engagement, which may in turn begin to impact effective evidence accumulation.

### Object recognition from memory (CBMT and CCMT)

#### Both studies: Lab and Online cohorts

Object recognition was assessed using the CBMT (lab cohort) and the CCMT (online cohort), paralleling the CFMT design. Due to ceiling effects in the CBMT, the online cohort completed the CCMT instead. Separate 2 × 3 ANOVAs (Group × Item Difficulty) showed main effects of Item Difficulty for both tasks: CBMT, *F*_Lab_(2, 106.9) = 22.6, *p*_Lab_ = <.001, *η^2^*_p_ = 0.3, BF_10_ > 1000, and CCMT, *F*_Online_(1.97, 146) = 144.83, *p*_Online_ <.001, *η^2^*_p_ = 0.66, BF_10_ > 1000. No group differences were observed in either task (all *p* ≥ .27, BF_10_ ≤ 0.44; see Supplementary Table 1, section 1.1; and Supplementary Figure 1, section 2.1).

### Face perception (CFPT)

#### Both studies: Lab and Online cohorts

Performance on the CFPT, which assesses face perception without memory demands, was indexed as the summed deviation between the response order and the target order (based on decreasing morph similarity). Separate 2 × 2 ANOVAs (Group × Orientation) in each cohort revealed strong main effects of Orientation (both *p* < .001, BF_10_ > 1000), but no Group differences (all *p* ≥ .52, BF_10_ ≤ 0.46; see Table 1).

**Table 1.**

|  |  | Imagers |  |  |  | Aphantasics |  |  |  |
| --- | --- | --- | --- | --- | --- | --- | --- | --- | --- |
|  |  | Lab |  | Online |  | Lab |  | Online |  |
|  |  | M | SD | M | SD | M | SD | M | SD |
| Orientation | Upright | 4.43 | 2.21 | 5.49 | 3.16 | 4.53 | 1.62 | 4.76 | 2.48 |
|  | Inverted | 7.33 | 1.85 | 7.89 | 2.03 | 7.59 | 2.12 | 7.95 | 1.98 |

*Note.* Means and standard deviations for mean deviations in upright vs. inverted face perception in the CFPT across groups and cohorts. Lower deviations indicate more precise perception of face similarities.

### Confidence ratings

Confidence was collected on a continuous scale from 0% (completely unsure) to 100% (completely sure) on each trial. As patterns were comparable across cohorts (Figure 4), analyses included Cohort as a factor.

**Figure 4.**
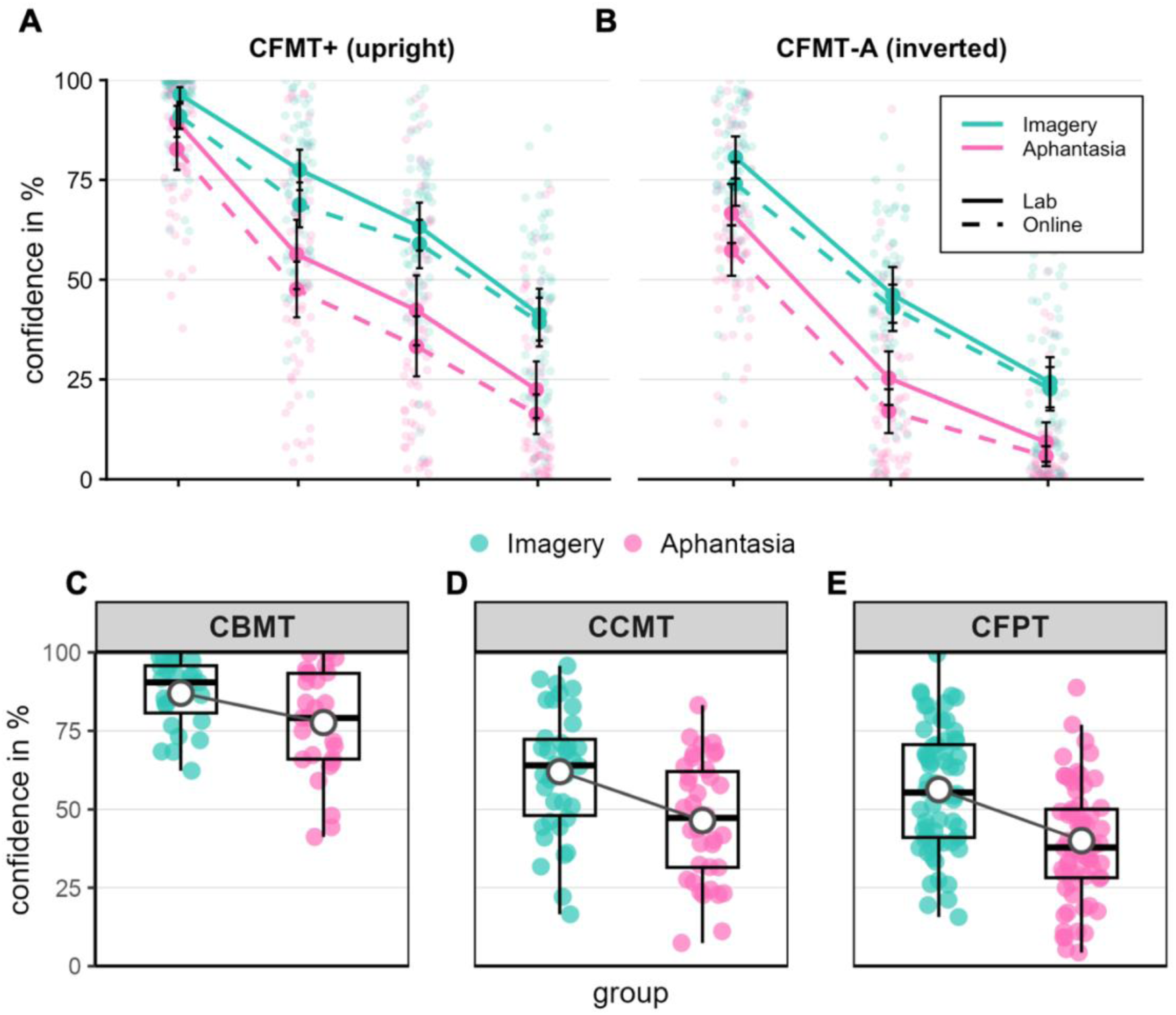
Confidence ratings. Top panel: mean confidence ratings given as percentages (0 = completely uncertain, 100 =completely certain) across test conditions for upright (CFMT+) (A) and inverted (CFMT-AM) (B) face recognition. Bottom panel: Mean confidence ratings by group (collapsed across cohorts and conditions) in object recognition of bicycles (CBMT; lap-participants only) and cars (CCMT; online participants only), and in face perception (CFPT; both cohorts). Colors (group) and line types (cohort): Turquoise = Typical Imagers, magenta = Aphantasics; solid line = Lab participants, dashed line = Online participants. Error bars signify standard errors of the mean (SE). Individual data points are shown with boxplots (medians = solid black lines; means = grey-edged white dots) and light grey lines connecting group means for visibility.

Because tasks used a 3AFC format, standard meta-d′ was not applicable. Instead, metacognition was assessed via confidence discrimination: participants’ ability to differentiate between their own correct and incorrect responses via confidence ratings. Due to low error rates, the same-images condition (level 1) was excluded.

#### Face recognition (CFMT+)

Confidence was analyzed using a 2 × 2 × 2 × 3 ANOVA (Cohort × Group × Accuracy × Item Difficulty). Participants were more confident in correct than incorrect responses (53.3% vs. 33%, *p* < .001, BF_10_ > 1000), and confidence decreased with increasing difficulty (*p* < .001, BF_10_ > 1000). This effect was qualified by an Accuracy × Item Difficulty interaction, *F*(2, 236.4) = 49.59, *p* < .001, *η^2^*_p_ = 0.30, BF_10_ > 1000, reflecting a steeper decline in confidence across levels of difficulty for correct trials (65.5%, 53.1% and 32%) than for incorrect trials (46.2%, 38.8% and 26.7%).

A main effect of group was observed, *F*(1, 118) = 38.31, *p* < .001, *η^2^*_p_ = 0.25, BF_10_ > 1000), with aphantasics reporting lower confidence than typical imagers across all difficulty levels (see Table 2). No effects involving Cohort were found (all *p* ≥ .75, all BF_10_ ≤ 0.36). Although a Group × Trial Accuracy × Item Difficulty interaction reached significance, *F*(2, 236) = 4.43, *p* = .013, η^2^_p_ = 0.04, it was not supported by Bayesian evidence (BF_10_ = 0.25) and is not considered further.

**Table 2.**

|  |  | Difficulty level |  |  |  |  |  |  |  |  |  |  |  |  |  |  |  |
| --- | --- | --- | --- | --- | --- | --- | --- | --- | --- | --- | --- | --- | --- | --- | --- | --- | --- |
|  |  | Same |  |  |  | Novel |  |  |  | Noise |  |  |  | Difficult |  |  |  |
| Test | Group | Imagers |  | Aphantasics |  | Imagers |  | Aphantasics |  | Imagers |  | Aphantasics |  | Imagers |  | Aphantasics |  |
|  |  | M | SD | M | SD | M | SD | M | SD | M | SD | M | SD | M | SD | M | SD |
| Test | CFMT+ | 93.2 | 9.6 | 85.6 | 16.1 | 72.1 | 18.3 | 51.3 | 25.4 | 60.6 | 20.1 | 37.1 | 26.6 | 40.1 | 20.7 | 18.9 | 19.3 |
|  | CFMT-AM | 76.7 | 17.9 | 61.2 | 22.1 | 44.2 | 20.3 | 20.6 | 19.5 | 23.6 | 18.5 | 7.3 | 11.5 |  |  |  |  |
|  | CBMT<sub>lab</sub> | 97.3 | 4.4 | 97.7 | 3.1 | 82.9 | 14 | 71 | 21.3 | 84.5 | 14.8 | 70.9 | 25.6 |  |  |  |  |
|  | CCMT<sub>online</sub> | 80.9 | 14 | 75.1 | 19.6 | 57.1 | 21.9 | 39.4 | 21 | 53.9 | 24.6 | 33.7 | 22.8 |  |  |  |  |

*Note.* Means and standard deviations of confidence ratings across memory tasks in percent (%), ranging from 0 to 100 with both cohorts collapsed.

Overall, both groups were similarly sensitive to accuracy and difficulty, but aphantasics reported consistently lower confidence (see Supplementary Figure 3, section 2.3.1). Similar reductions were observed for inverted faces and object recognition (Figure 4).

#### Face perception (CFPT)

To test whether reduced confidence extends beyond imagery-based memory tasks,, we examined confidence in face perception. This is important because aphantasia is most often defined via the VVIQ, which depends on imagery vividness; thus, perceptual tasks that do not overtly rely on imagery allow us to assess whether reduced confidence generalizes beyond imagery-dependent processes.

As CFPT responses yield continuous deviation scores, metacognitive sensitivity could not be computed. Thus, absolute confidence was analyzed using a 2 × 2 × 2 ANOVA (Group × Cohort × Orientation: upright-inverted). Confidence was lower for inverted than upright faces, (*M* = 55.2% vs. 41.4%, *F*(1, 126) = 121.63, *p* < .001, *η^2^*_p_ = 0.49, BF_10_ > 1000). There was also a main effect of Group, *F*(1, 126) = 20.93, *p* = 1.12×10^−5^, *η^2^*_p_ = 0.14, BF_10_ > 1000, with aphantasics reporting lower confidence than typical imagers for both upright (47.2% vs. 63.5%) and inverted faces (32.9% vs. 49.5%; both *p* < .001, *d* ≥ 0.78, BF_10_ > 1000) despite comparable performance. No main effect of Cohort was observed (*p* = .99, BF_10_ = 2).

A Cohort × Orientation interaction emerged, *F*(1, 126) = 8.57, *p* = .004, *η^2^*_p_ = 0.06, BF_10_ = 7.29, but follow-up comparisons were not significant (both *p* = .63, BF_10_ < 0.3).

### Meta-analysis of group differences in short form CFMT: Aphantasics vs. typical imagers

To our knowledge, four studies, including the present one, have compared aphantasics and typical imagers using the standardized CFMT (short form), enabling a meta-analytic assessment of group differences.

Across studies, aphantasics showed reliably lower face recognition performance at all difficulty levels: same-images, *μ̂*_same_ = −0.248, 95% CI: [−0.439, −0.057], z = −2.55, *p* = .011; novel-images, *μ̂*_novel_ = −0.408, 95% CI: [−0.616, −0.201], z = −3.86, *p* < .001; noisy-images, *μ̂*_noise_ = −0.308, 95% CI: [−0.499, −0.118], z = −3.17, *p* = 0.0015 (Figure 5). However, note that effect sizes at the first level of difficulty (i.e., in the same-images condition) should be interpreted with caution, as ceiling effects restrict variance and can inflate standardized estimates. There was no evidence of heterogeneity across samples (all *Q*(4) ≥ 5.69, all *p* ≥ .337), variation in effect sizes across studies was not greater than expected by chance. Notably, the results were unchanged when restricting analyses to participants with floor-level VVIQ scores (VVIQ = 16).

**Figure 5.**
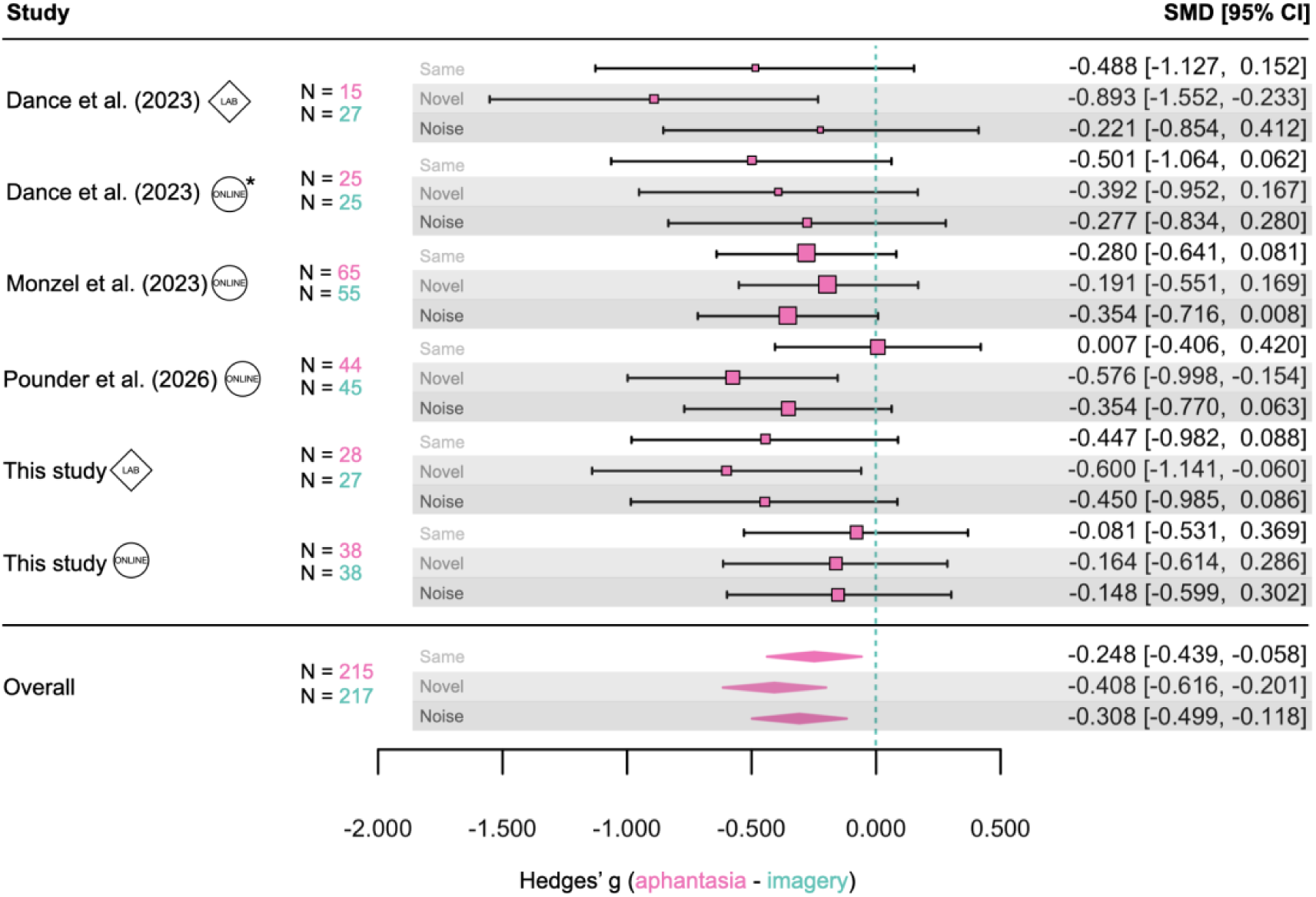
Forest plot showing three independent meta-analyses for each difficulty level of the short form CFMT (i.e., same- images, novel images, and images with noise). Each meta-analysis includes data from three studies separated by lab- vs. online cohorts. *Online data in Dance et al. (2023) were collected under supervision of an experimenter in a joint Skype call.

### Questionnaires

Consistent with previous studies (Dance et al., 2021; Dawes et al., 2020), aphantasics and typical imagers differed on all PSI-Q subscales (all *p* < .001) and on the object subscale of the OSIVQ (*p* < .001), but not on the spatial or verbal subscales (both *p* = .35). Significant group differences were also observed on all SAM subscales (episodic, future, and semantic memory: all *p* < .001; spatial memory: *p* = .01). Full statistics are reported in Supplementary Table 6, section 1.2.

Following the experimental tasks, participants provided a global assessment of their perceived performance on each task using a scale ranging from 0 to 100. Consistent with the trial-by-trial confidence ratings, aphantasics reported lower perceived performance than typical imagers on all tasks, although not in the Bikes/Cars task (all *p* ≤ .01; Bikes/Cars: *p* = 0.72; see Supplementary Figure 6, section 2.4.2).

Participants also completed a post-experimental questionnaire assessing imagery use in everyday life, dream experiences, and related phenomenological characteristics (Supplementary Figure 5, section 2.4.2).

Finally, participants were asked to describe the strategy they had used to remember faces during the experiment. Responses were initially coded as relying primarily on specific facial features, processing the face as a whole, or using both approaches. For the statistical analysis, the latter two categories were combined into a single holistic-processing category. A chi-square test of independence revealed a significant association between group and reported strategy, χ^2^(1, N = 101) = 6.87, *p* = .0088, BF_10_ = 5.65. Aphantasics more frequently reported relying on specific facial features, whereas typical imagers were more likely to report either processing the face as a whole or using a combination of whole-face and feature-based strategies (Supplementary Figure 5, bottom right panel, section 2.4.2).

## Discussion

Our findings clarify and extend current accounts of how visual imagery contributes to face recognition. Consistent with previous findings (Monzel et al., 2023b; Dance et al., 2023), individuals with typical imagery ability outperformed aphantasics on the CFMT, a standardized face recognition task. In the present data, this advantage manifested as higher accuracy in the lab cohort and more efficient evidence accumulation (drift rates) across both lab and online samples. A meta-analysis combining four studies (N = 432), further confirmed a reliable, albeit modest, advantage for typical imagers across all difficulty levels of the short form CFMT, i.e., same-, novel-, and noisy face images. Importantly, in our study, group differences were most pronounced at the highest and previously untested difficulty level of the long form CFMT (CFMT+; Russell et al., 2009), suggesting that imagery-related benefits emerge most clearly under demanding perceptual and mnemonic conditions.

One possible account for the better performance of the imagery group is that visual imagery supports the maintenance or retrieval of more detailed visual representations of previously encountered faces. This interpretation is broadly consistent with evidence that visual imagery recruits neural representations that overlap with those involved in visual perception (for a review, see Pearson, 2019). Individuals with stronger imagery may therefore be better able to reconstruct facial information when comparing stored representations with currently perceived faces, particularly under challenging recognition conditions. Importantly, this does not imply that aphantasics are unable to encode or maintain visual information per se (see Weber et al., 2024). Rather the difference might lie in the format or quality of the representations used during retrieval. In this sense, imagery may facilitate more effective top-down reinstatement of perceptual information (Ortiz-Tudela et al., 2023), which could support recognition across changes in viewpoint, lighting, or visual noise (Wallace, 1988). Consistent with this view, mnemonic reinstatement in early visual areas (V1 and V2) depends on whether retrieval is guided via episodic or semantic routes (Ortiz-Tudela et al., 2023), with verbal strategies potentially providing less precise visual content carried in high-level feedback processes (Bergmann & Ortiz-Tudela, 2023). This may explain why group differences became most pronounced in the CFMT+, where successful recognition likely depends on maintaining stable yet flexible face representations under highly demanding perceptual conditions.

Face recognition relies on both visual and verbal processing routes (Clark & Paivio, 1987). Visual encoding involves multiple mechanisms, including holistic (processing the face as a unified whole), configural (encoding relational information between facial features) and featural processing (encoding facial features; Farah et al., 1986; Maurer et al., 2002). Featural processing can be supported by visually derived semantic labels (Bruce & Young, 1986), such as “has a wide jaw line”, or broader associations, such as “looks like my uncle”, both of which can aid recognition (Klatzky et al., 1982; McKelvie, 1976). However, optimal face recognition is typically associated with holistic processing, in which facial features and their spatial relations are integrated into a unified representation (Richler et al., 2011; Yin, 1969). Consistent with previous studies, participants with aphantasia reported a stronger reliance on featural strategies, whereas imagers integrate both, featural and holistic information (see Supplementary Figure 5). This pattern aligns with evidence that aphantasics preferentially use verbal labeling and feature-based encoding across visual memory tasks (Bainbridge et al., 2021; Monzel et al., 2024; Keogh & Pearson, 2021, but see Reeder where other strategies are reported).

Consistent with this interpretation, findings from the verbal overshadowing literature may also be relevant to understanding face processing in aphantasia. Verbal overshadowing refers to the impairment in later recognition that can occur after verbally describing a previously encountered face (Schooler & Engstler-Schooler, 1990; Schooler, 2002). Although verbal descriptions can sometimes support recognition, verbalization may impair performance when they promote feature-based analysis at the expense of holistic face processing. Interestingly, recent evidence suggests that aphantasics do not show the typical verbal overshadowing effect, although the study did not specifically examine face recognition (Monzel et al., 2024). One possible interpretation is that aphantasics already rely relatively more on feature-based strategies during recognition, such that verbalization produces little additional shift in processing strategy and therefore has a reduced impact on subsequent recognition.

An interpretation along such lines may help explain mixed results across studies with differing methodologies. For example, tasks that include additional external features (e.g., haircuts or accessories; McKelvie, 1994; Milton et al., 2021), may promote the use of verbally mediated strategies by increasing the availability of diagnostic features^3^. Such features could stabilize verbal codes across encoding and retrieval, allowing participants to compensate for weaker visual representations. A similar pattern emerges in the visual working memory literature: aphantasics and typical imagers often show comparable accuracy (Knight et al., 2022; Pounder et al., 2022; Weber et al., 2024), but aphantasics tend to respond slower (Liu & Bartholomeo, 2023; Pounder et al., 2022), potentially reflecting greater reliance on non-visual strategies (Kay et al., 2024; Reeder et al., 2024). Crucially, when task demands emphasize high-precision processing of visual information, imagery ability appears to confer an advantage (Jacobs et al., 2018). In this context, the CFMT+, a test designed to identify super-recognizers (Russell et al., 2009), likely places particularly strong demands on memory precision.

We also examined performance for inverted faces. Consistent with the well-established face inversion effect (Yin, 1969), recognition accuracy declined in both groups. However, in the lab cohort, group differences were more pronounced for inverted than upright faces and were evident across all difficulty levels. This could indicate that aphantasics are more strongly affected when holistic processing is disrupted. Two, not mutually exclusive, explanations may account for this pattern. First, typical imagers may rely more effectively on holistic processing overall, allowing them to better sustain performance even under inversion (Wallace, 1988). Second, imagers may also benefit from more efficient featural processing. Although aphantasics reported primarily feature-based strategies, typical imagers appeared to combine featural and holistic information (see Supplementary Figure 5, section 2.4.2). While one might expect reduced group differences when holistic information is disrupted, evidence suggests that featural processing itself can benefit from visual imagery (Lobmaier & Mast, 2008) and may support superior recognition performance even when forced to adopt feature-based sampling strategies (Dunn et al., 2022). This raises the possibility that, even under conditions where holistic processing is constrained, typical imagers may retain an advantage through more effective use of both holistic and featural information.

Notably, McKelvie (1994) did not find differences between “good” and “poor” visualizers in inverted face recognition. This discrepancy may reflect key methodological differences: McKelvie’s study did not include individuals with markedly reduced or absent imagery, limiting the range of imagery abilities, and stimuli included external features, which are known to reduce group differences between prosopagnosics and controls (Duchaine & Weidenfeld, 2003). One important limitation in our study is that the CFMT-AM uses a different set of target faces to the CFMT, which prevented a direct within-subject comparison of upright and inverted performance. Future studies should examine inversion effects using designs that allow more controlled comparisons across conditions.

In contrast to prior findings (Monzel et al., 2023b), we observed no group differences in object recognition for either cohort. However, these results should be interpreted cautiously: the CBMT showed ceiling effects in the lab sample (accuracy > 85%), and the lack of differences in CCMT performance in the online cohort might be influenced by reduced response caution in typical imagers, as observed in the other tasks. Accordingly, the absence of group differences in object recognition should be considered preliminary.

Our study also assessed face perception using the CFPT, which requires participants to rank face morphs by similarity to a target. Despite prior evidence linking CFPT performance to face memory (Fysh & Bindemann, 2018), we found no group differences between aphantasics and typical imagers. Importantly, unlike earlier findings (e.g., Dance et al., 2023), we assessed face perception and memory within the same individuals, allowing a more direct comparison of perceptual and mnemonic processing. Our results therefore provide stronger evidence for a dissociation between face memory and face perception in aphantasia that is not attributable to differences in task demands or participant samples. More broadly, group differences appear to emerge primarily when visual information must be internally maintained and matched to a target, whereas no differences are observed when all information is externally available, as in the CFPT. This interpretation appears to contrast with recent findings of Pounder et al. (2026), who reported impairments in both face memory and face perception using the Oxford Face Matching Test (OFMT; Stantic et al., 2021). However, unlike the CFPT, which allows continuous comparison of faces (mean exposure in the present study = 41.52 s), the OFMT presents face pairs for only 1600 ms before collecting similarity and identity judgments. Consequently, although the OFMT derives a refined measure of face perception from similarity ratings, these judgments are made after stimulus offset and may therefore rely more heavily on internally maintained visual representations than those made while the faces remain visible. This difference may explain why group differences emerged in the OFMT but not in the CFPT and is consistent with our finding that impairments arise primarily when visual information must be internally maintained. These findings argue against accounts proposing a global face-processing deficit in aphantasia (e.g., Maw et al., 2025) and instead support an imagery-specific impairment. Although lesion-acquired aphantasia has been associated with damage to face-processing regions (including the proposed fusiform imagery node) and with prosopagnosia-like symptoms (Kutschke et al., 2026), evidence from non-lesion samples relies largely on self-reported face-recognition difficulties (e.g., Dance et al., 2023; Zeman et al., 2020). In contrast, behavioral studies, including the present findings, provide little evidence for a substantial impairment in face recognition associated with aphantasia (Monzel et al., 2023b).

We also assessed trial-by-trial decision confidence and found consistent group differences: typical imagers reported higher confidence than aphantasics. Notably, this effect generalized across stimulus categories (faces, bicycles, cars) and task types and was observed even in the absence of accuracy differences in the face perception task.

Previous work highlights different aspects of this pattern. Liu & Bartolomeo (2023) emphasize stimulus type, reporting reduced confidence for shapes and faces, but not for colors, words, or spatial arrangements. In contrast, Wittmann & Şatırer (2022) focus on task demands, suggesting that confidence differences emerge in contexts requiring vivid recollection, regardless of modality. Our results extend this perspective by suggesting that reduced confidence may arise broadly whenever tasks might be associated with imagery. This interpretation is consistent with the *self-stigmatization hypothesis*, whereby aphantasics may anticipate poorer performance in imagery-related contexts (Monzel et al., 2024). Such expectations may be reinforced by broader self-perceptions: e.g., aphantasics often report feeling “different” from others (Mawtus et al., 2024). Notably, in all the above studies on confidence, including our own, aphantasic participants were recruited through dedicated databases or imagery-focused forums, potentially reinforcing such expectations. The use of a continuous confidence scale may have further amplified these differences by capturing finer-grained variability than traditional 4- or 5-point rating scales (Norman & Price, 2015).

A limitation of the present study is that the 3AFC design precluded the use of formal metacognitive measures such as meta-d’ (Maniscalco & Lau, 2012). Instead, we approximated metacognitive sensitivity by comparing confidence ratings for correct versus incorrect trials when sufficient errors were available. This analysis revealed no group differences, suggesting that the observed confidence gap reflects differences in confidence bias, rather than metacognitive sensitivity. Nonetheless, the absence of signal-detection-based measures remains a methodological limitation.

Finally, our study highlights important challenges in conducting online research with low-prevalence populations such as aphantasia. Given its rarity (1.5 – 4%; Beran et al., 2024; Dance et al., 2022), much of the literature relies on online recruitment and testing (e.g., Kay et al., 2024; Monzel et al., 2023b; Speed et al., 2025; Wittmann & Şatırer, 2022), yet our findings show that online and lab data can diverge in meaningful ways. While typical imagers outperformed aphantasics at the highest level of the CFMT+ across both cohorts, and in the inverted task, several group differences observed in the lab were attenuated or absent online.

Drift diffusion modelling revealed a distinct response profile in online typical imagers, marked by faster responding and reduced caution (see also Kay et al., 2024, for a similar finding from a mental rotation task), indicating differences in decision strategies and likely task engagement. Such discrepancies are difficult to control in remote settings and may be particularly pronounced in aphantasia research: aphantasic participants, often recruited from dedicated communities, may be highly motivated, whereas online control participants, often recruited from crowdsourcing platforms (e.g., Prolific, Amazon Mechanical Turk) may feel less connected to the research and might therefore be less engaged. The absence of in-person supervision may further amplify these discrepancies. Notably, several online studies have reported null findings (e.g., Delhaye et al., 2026; Liu & Bartolomeo, 2023; Pounder et al., 2024) which our results suggest may partly reflect variability in engagement, motivation, or perceived task relevance rather than a true absence of group differences.

This line of observation underscores the need for careful methodological control in online studies. Future work may benefit from (1) prescreening imagery ability within general populations to reduce self-selection biases (e.g., in Cabbai et al., 2024; Weber et al., 2024); (2) directly measuring and equating task engagement (e.g., via performance-contingent incentives, explicit effort checks, or post-task motivation ratings; and (3) incorporating hybrid or monitored testing procedures to improve compliance and standardization (e.g., in part by Dance et al., 2023, Reeder et al., 2024 and Azañón et al., 2024).

In sum, our findings show that visual imagery ability is associated with a reliable advantage in face recognition under high perceptual and mnemonic demands. This advantage was most pronounced at the highest difficulty level of the CFMT+. A meta-analysis across four studies further indicates a small but consistent advantage for typical imagers across standard CFMT conditions, suggesting that imagery-related differences are stable but do not seem to increase monotonically with successive levels of task difficulty. More broadly, our results point to the importance of task demands and testing context in revealing imagery-related effects. In addition, the consistent confidence differences observed here suggest systematic differences in self-evaluation among aphantasics that extends beyond memory-specific tasks. Together, these findings refine our understanding of how visual imagery contributes to face recognition and highlight key methodological considerations for future research.

## Acknowledgements

We thank Nadine Schröder for helping with collecting the laboratory data and Bradley Duchaine for providing test materials.

## Footnotes

1 Note, that unlike the facilitatory effect reported earlier, adding external features (e.g., haircut) at test changes the retrieval context and may impair recognition, particularly for unfamiliar faces (Bruce, 1982).

2 Note that the authors did not perform analyses for each separate cohort in the original publication (see Dance et al., 2023).

3 This seems especially the case for the *Famous Face Test*, where stimuli are familiar faces of public figures (Milton et al., 2021).

