## Supplementary Materials for "The Role of Visual Imagery in Face Recognition and Confidence Revisited: New Evidence from Aphantasia, Sampling Context Effects, and a Meta-Analysis"

#### 1 Additional Tables

##### 1.1 Memory performance

Table 1: Accuracy

| | | Difficulty level | | | | | | | | | | | | | | | |
|---|---|---|---|---|---|---|---|---|---|---|---|---|---|---|---|---|---|
| | | Same | | | | Novel | | | | Noise | | | | Difficult | | | |
| | **Group** | Imagery | | Aphantasia | | Imagery | | Aphantasia | | Imagery | | Aphantasia | | Imagery | | Aphantasia | |
| | **Cohort** | Lab | Online | Lab | Online | Lab | Online | Lab | Online | Lab | Online | Lab | Online | Lab | Online | Lab | Online |
| | | **M** | **M** | **M** | **M** | **M** | **M** | **M** | **M** | **M** | **M** | **M** | **M** | **M** | **M** | **M** | **M** |
| | | *SD* | *SD* | *SD* | *SD* | *SD* | *SD* | *SD* | *SD* | *SD* | *SD* | *SD* | *SD* | *SD* | *SD* | *SD* | *SD* |
| **Test** | CFMT+ | **99.2** | **95.8** | **97.6** | **95.2** | **84.2** | **75.4** | **75.1** | **72.7** | **74.4** | **66.6** | **65.8** | **63.6** | **54.9** | **46** | **41** | **43** |
| | | *2.5* | *6.9* | *4.1* | *7.4* | *12.2* | *17.8* | *17.1* | *14.9* | *17.3* | *19* | *20.3* | *20.5* | *14.3* | *13.1* | *13.6* | *13.8* |
| | CFMT-AM | **95.1** | **81.4** | **87.3** | **84.5** | **56.1** | **45.8** | **47** | **45.7** | **39.5** | **34.5** | **32.7** | **34.5** | | | | |
| | | *6.4* | *18.8* | *11.2* | *12.9* | *13.1* | *12.6* | *10.3* | *13.3* | *9.8* | *9.8* | *7.4* | *10.4* | | | | |
| | CBMT | **97.5** | | **97** | | **86.5** | | **85.2** | | **92.9** | | **86.3** | | | | | |
| | | *3.9* | | *5.8* | | *14* | | *11.7* | | *9.2* | | *19.9* | | | | | |
| | CCMT | | **86.1** | | **88.6** | | **59.2** | | **62** | | **66.5** | | **69.9** | | | | |
| | | | *11.1* | | *9.9* | | *12.5* | | *15.3* | | *16.7* | | *17.3* | | | | |

*Note*. Means and standard deviations of accuracy across memory tasks in percent correct (%). Note that object recognition tests were split between cohorts, i.e., lab participants underwent the CBMT, whereas online participants performed the CCMT.

Table 2: Response times

| | | Difficulty level | | | | | | | | | | | | | | | |
|---|---|---|---|---|---|---|---|---|---|---|---|---|---|---|---|---|---|
| | | Same | | | | Novel | | | | Noise | | | | Difficult | | | |
| | **Group** | Imagery | | Aphantasia | | Imagery | | Aphantasia | | Imagery | | Aphantasia | | Imagery | | Aphantasia | |
| | **Cohort** | Lab | Online | Lab | Online | Lab | Online | Lab | Online | Lab | Online | Lab | Online | Lab | Online | Lab | Online |
| | | **M** | **M** | **M** | **M** | **M** | **M** | **M** | **M** | **M** | **M** | **M** | **M** | **M** | **M** | **M** | **M** |
| | | *SD* | *SD* | *SD* | *SD* | *SD* | *SD* | *SD* | *SD* | *SD* | *SD* | *SD* | *SD* | *SD* | *SD* | *SD* | *SD* |
| **Test** | CFMT+ | **3.12** | **4.17** | **3.54** | **4.18** | **6.72** | **4.43** | **6.36** | **6.32** | **6.38** | **3.95** | **6.24** | **5.36** | **8.2** | **4.73** | **6.78** | **6.13** |
| | | *1.09* | *1.27* | *1.25* | *1.47* | *4.76* | *1.75* | *3.22* | *2.76* | *3.87* | *1.3* | *2.79* | *2.27* | *4.45* | *1.89* | *3.51* | *2.54* |
| | CFMT-AM | **4.63** | **3.62** | **4.63** | **4.94** | **7.76** | **4.29** | **6.48** | **5.85** | **6.49** | **3.44** | **4.91** | **4.51** | | | | |
| | | *1.37* | *1.5* | *1.62* | *1.52* | *3.78* | *1.95* | *2.47* | *2.97* | *3.45* | *1.71* | *2.41* | *2.58* | | | | |
| | CBMT | **2.98** | | **2.59** | | **5.92** | | **6.16** | | **4.63** | | **4.59** | | | | | |
| | | *1.22* | | *0.9* | | *2.71* | | *2.65* | | *1.75* | | *2.08* | | | | | |
| | CCMT | | **4.44** | | **5.55** | | **6.08** | | **7.72** | | **4.6** | | **6.23** | | | | |
| | | | *1.55* | | *1.99* | | *2.77* | | *2.85* | | *1.98* | | *2.26* | | | | |

*Note*. Means and standard deviations of participant's median response times in seconds (s). Note that object recognition tests were split between cohorts, i.e., lab participants underwent the CBMT, whereas online participants performed the CCMT.

Table 3: Boundary separation

| | | Difficulty level | | | | | | | | | | | | | | | |
|---|---|---|---|---|---|---|---|---|---|---|---|---|---|---|---|---|---|
| | | Same | | | | Novel | | | | Noise | | | | Difficult | | | |
| | **Group** | Imagery | | Aphantasia | | Imagery | | Aphantasia | | Imagery | | Aphantasia | | Imagery | | Aphantasia | |
| | **Cohort** | Lab | Online | Lab | Online | Lab | Online | Lab | Online | Lab | Online | Lab | Online | Lab | Online | Lab | Online |
| | | **M** | **M** | **M** | **M** | **M** | **M** | **M** | **M** | **M** | **M** | **M** | **M** | **M** | **M** | **M** | **M** |
| | | *SD* | *SD* | *SD* | *SD* | *SD* | *SD* | *SD* | *SD* | *SD* | *SD* | *SD* | *SD* | *SD* | *SD* | *SD* | *SD* |
| **Test** | CFMT+ | **4.05** | **4.07** | **4.17** | **4.59** | **5.37** | **4.22** | **5.22** | **5.01** | **4.88** | **3.64** | **4.76** | **4.34** | **5.19** | **3.88** | **4.76** | **4.48** |
| | | *0.79* | *0.84* | *0.74* | *0.97* | *1.27* | *1.01* | *1.1* | *1.23* | *1.06* | *0.78* | *1.11* | *1.13* | *1.16* | *0.9* | *1.78* | *1.06* |
| | CFMT-AM | **4.95** | **3.86** | **4.59** | **4.65** | **4.6** | **3.34** | **4.07** | **3.73** | **4.28** | **3.01** | **3.64** | **3.48** | | | | |
| | | *0.89* | *1.16* | *0.96* | *0.97* | *1.15* | *0.91* | *0.89* | *1.05* | *1.15* | *0.87* | *0.96* | *1.09* | | | | |

*Note*. Means and standard deviations of the estimated boundary separation parameter.

Table 4: Drift rates

| | | Difficulty level | | | | | | | | | | | | | | | |
|---|---|---|---|---|---|---|---|---|---|---|---|---|---|---|---|---|---|
| | | Same | | | | Novel | | | | Noise | | | | Difficult | | | |
| | **Group** | Imagery | | Aphantasia | | Imagery | | Aphantasia | | Imagery | | Aphantasia | | Imagery | | Aphantasia | |
| | **Cohort** | Lab | Online | Lab | Online | Lab | Online | Lab | Online | Lab | Online | Lab | Online | Lab | Online | Lab | Online |
| | | **M** *SD* | **M** *SD* | **M** *SD* | **M** *SD* | **M** *SD* | **M** *SD* | **M** *SD* | **M** *SD* | **M** *SD* | **M** *SD* | **M** *SD* | **M** *SD* | **M** *SD* | **M** *SD* | **M** *SD* | **M** *SD* |
| **Test** | CFMT+ | **1.21** *0.18* | **1.17** *0.19* | **1.04** *0.17* | **0.94** *0.19* | **0.57** *0.24* | **0.5** *0.32* | **0.41** *0.25* | **0.36** *0.22* | **0.4** *0.25* | **0.32** *0.26* | **0.25** *0.24* | **0.23** *0.26* | **0.07** *0.15* | **-0.04** *0.15* | **-0.11** *0.16* | **-0.1** *0.16* |
| | CFMT-AM | **0.69** *0.05* | **0.66** *0.08* | **0.61** *0.08* | **0.6** *0.07* | **0.05** *0.09* | **-0.01** *0.08* | **-0.03** *0.07* | **-0.04** *0.09* | **-0.15** *0.06* | **-0.2** *0.06* | **-0.23** *0.05* | **-0.22** *0.05* | | | | |

*Note*. Means and standard deviations of the estimated drift rate parameter across memory tasks.

Table 5: CFMT short form ANOVA results on accuracy

| **Effect** | **Statistics** | ***p*-value** | **Effect size ($\eta^2_p$)** | **$BF_{10}$** |
|---|---|---|---|---|
| Item Difficulty | $F(1.72, 91.2) = 107.9$ | < .001 | 0.67 | > 1000 |
| Group | $F(1, 53) = 4.48$ | .039 | 0.08 | 2.06 |
| Group x Item Difficulty | $F(1.72, 91.2) = 2.31$ | .11 | 0.04 | 1.45 |

*Note*. Lab performance (accuracy) in the short-form version of the CFMT. See full statistics in Table 1, including difficulty levels: same, novel and noise.

##### 1.2 Questionnaires

Table 6: Questionnaires

| Test | Subscale | Statistics | *p*-value | Effect size (d) | $BF_{10}$ |
|---|---|---|---|---|---|
| VVIQ | | *t*(1, 128) = 26.29 | < .001*** | 4.61 | > 1000 |
| PSI-Q | | *t*(1, 128) = 18.78 | < .001*** | 3.3 | > 1000 |
| | *appearance* | *t*(1, 128) = 26.62 | < .001*** | 4.67 | > 1000 |
| | *sound* | *t*(1, 128) = 12.99 | < .001*** | 2.28 | > 1000 |
| | *smell* | *t*(1, 128) = 17.8 | < .001*** | 3.12 | > 1000 |
| | *taste* | *t*(1, 128) = 17.69 | < .001*** | 3.1 | > 1000 |
| | *touch* | *t*(1, 128) = 14.11 | < .001*** | 2.48 | > 1000 |
| | *bodily sensation* | *t*(1, 128) = 12.26 | < .001*** | 2.15 | > 1000 |
| | *feeling (emotion)* | *t*(1, 128) = 9.2 | < .001*** | 1.61 | > 1000 |
| OSIVQ | | *t*(1, 110) = 7.75 | < .001*** | 1.47 | > 1000 |
| | *object* | *t*(1, 110) = 17.79 | < .001*** | 3.36 | > 1000 |
| | *spatial* | *t*(1, 110) = -1.36 | .35 | -0.26 | 0.46 |
| | *verbal* | *t*(1, 110) = -1.22 | .35 | -0.23 | 0.39 |
| SAM | | *t*(1, 111) = 11.38 | < .001*** | 2.14 | > 1000 |
| | *episodic* | *t*(1, 111) = 8.08 | < .001*** | 1.52 | > 1000 |
| | *facts* | *t*(1, 111) = 3.99 | < .001*** | 0.75 | 186.75 |
| | *spatial* | *t*(1, 111) = 3.01 | .01* | 0.57 | 10.57 |
| | *future* | *t*(1, 111) = 13.92 | < .001*** | 2.62 | > 1000 |

*Note*. Summary of independent samples t-tests on questionnaire scores, Holm-corrected for multiple comparisons. Asterisks indicate significance levels: *: $p < .05$, **: $p < .01$, ***: $p < .001$.

#### 2 Additional Figures

##### 2.1 Object recognition: Cambridge Bike Memory Test / Cambridge Car Memory Test

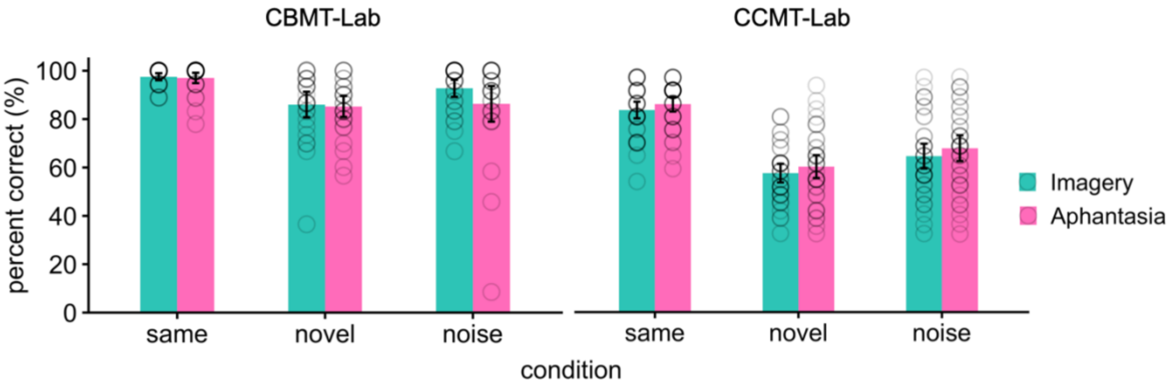


*Figure 1*. Mean object recognition accuracy as percentages of correct answers (3-AFC) for each level of difficulty, i.e., same, novel, and noisy images. Error bars represent 95% CIs. Note that only lab-participants underwent the CBMT due to consistent high-level performance. Hence, online-participants were given the CCMT to test for object recognition.

##### 2.2 Face perception: Cambridge Face Perception Test

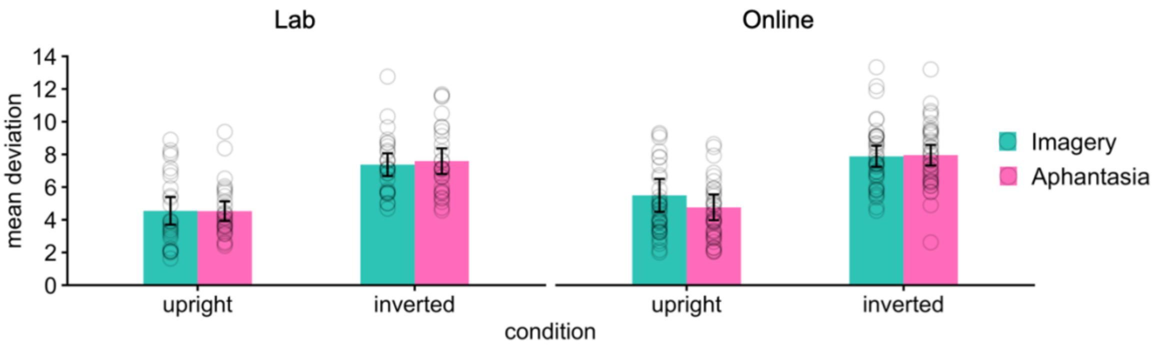


*Figure 2*. Mean deviation of target vs. selected face orders within a set of six faces for two different orientations, i.e., upright vs. inverted.

##### 2.3 Memory Confidence

2.3.1 Cambridge Face Memory Test: Correct vs. incorrect trials

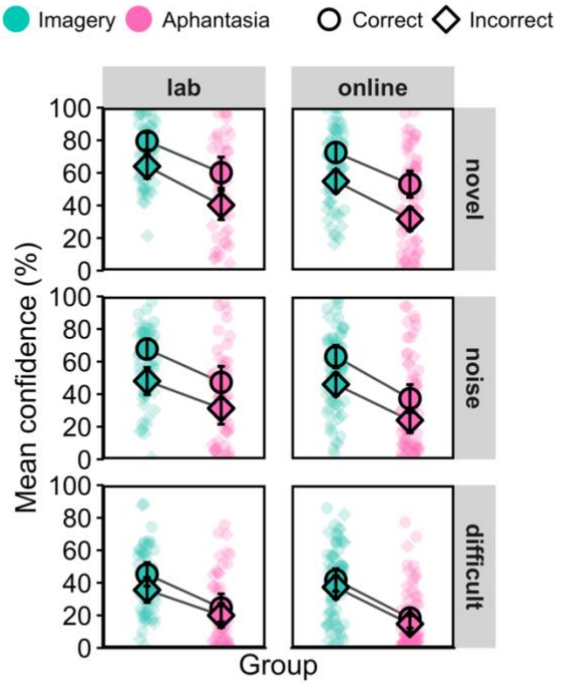


*Figure 3*. Cohort by group mean confidence ratings split by correct vs. incorrect trials in 3 levels of difficulty, i.e., novel, noisy, and difficult images in the CFMT+. Note that the first level, same images, was excluded from this analysis due to too few incorrect trials. Asterisks represent significance levels: *: $p < .05$, **: $p < .01$, ***: $p < .001$.

##### 2.4 Questionnaires

2.4.1 Survey of autobiographical memories (SAM; Palombo et al., 2013)

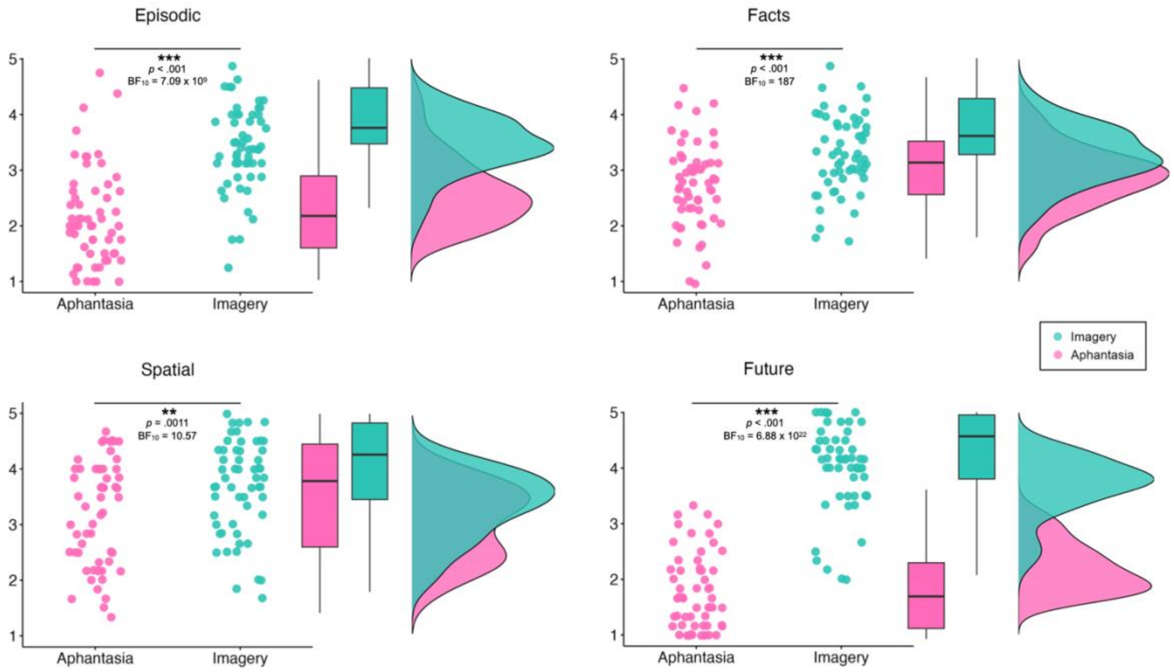


*Figure 4*. Raincloud plots representing mean values scored on each subscale of the SAM (i.e., episodic memory, memory of facts, spatial memory and memories about the future) by group (based on 58 aphantasics and 55 imagers). Asterisks represent significance levels: *: $p < .05$, **: $p < .01$, ***: $p < .001$.

2.4.2 Post experimental questionnaire

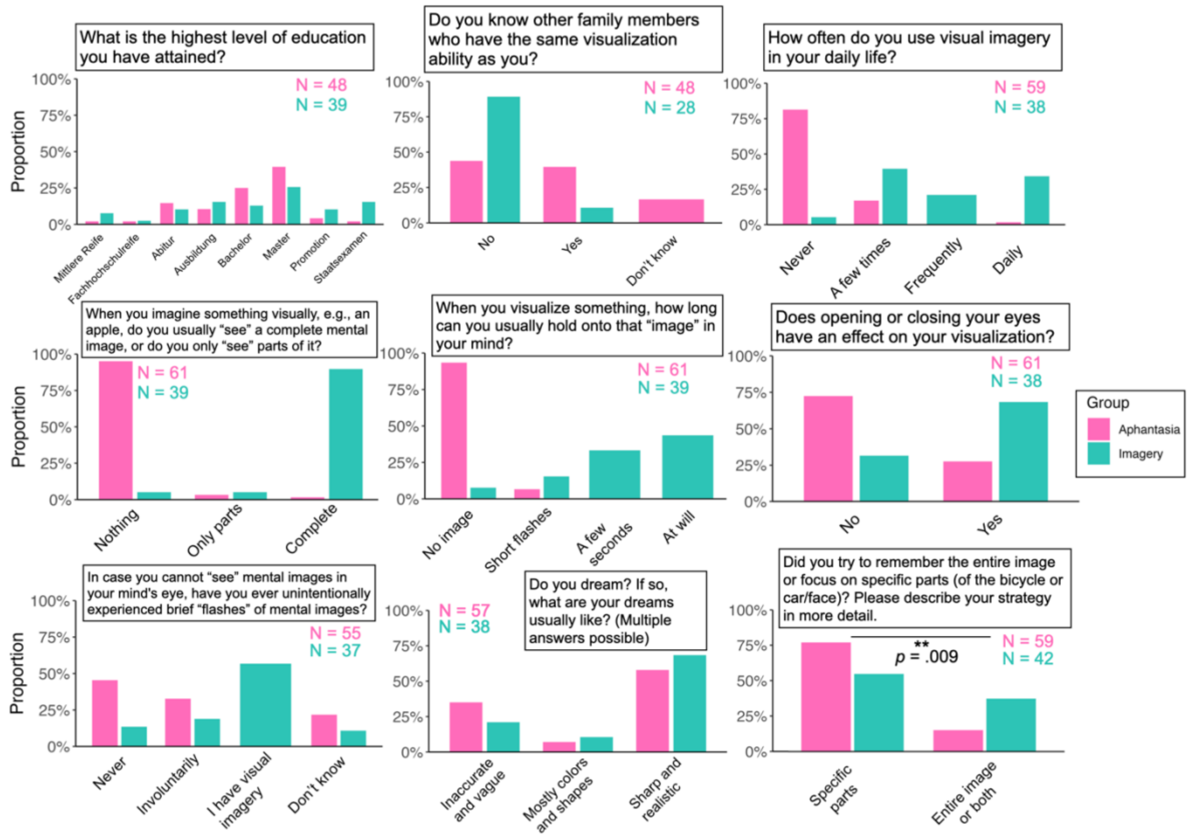


*Figure 5*. Bar plots representing the frequency of answers to the post-experimental questionnaire including strategy use (bottom right panel). Asterisks represent significance levels: *: p < .05, **: p < .01, ***: p < .001, n.s.: not significant. Note, that group comparisons were only performed on strategy use, i.e., the bottom right panel.

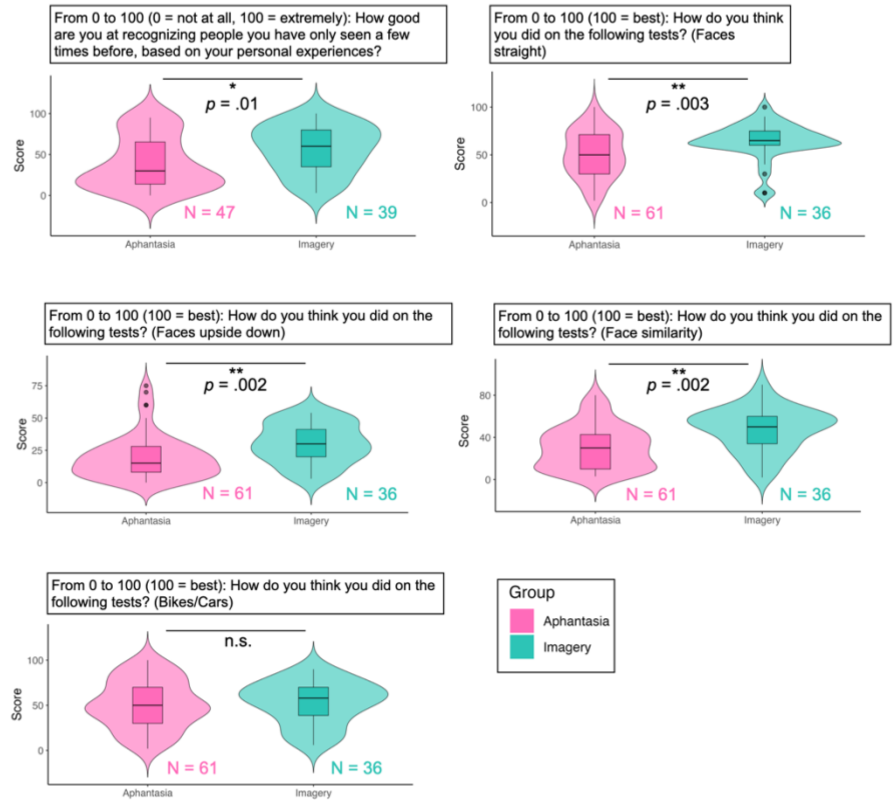


*Figure 6*. Violin box plots representing mean confidence scores based on general post-experimental assessments. Asterisks represent significance levels: *: $p < .05$, **: $p < .01$, ***: $p < .001$, n.s.: not significant.

### 3 Additional Methods

#### 3.1 Stimuli

In the CFMT+, study face images measured approximately 180 × 220 pixels, subtending approximately 3.9° (width) × 4.8° (height) of visual angle. During test trials, three faces were presented simultaneously with the middle face centered on the screen and the left and right faces spaced approximately 5.7° apart to either side (center-to-center). In the review phase, prior to the novel- and noisy-images levels, six faces (2 rows × 3 columns) were shown in a 700 × 700-pixel image centered on screen. The absolute sizes of the faces were consistent across study, test and review phases. Bicycle images in the CBMT were displayed at approximately 6.9° × 6.9° degrees of visual angle and were shown side-by-side at the center of the screen during test trials. In the CFPT (lab version), the target face was displayed at the top center of the screen (13.6° of visual angle from the centroid position) subtending 6.5° × 6.5° of visual angle. The six response faces were arranged in a horizontal row, each subtending 6.5° × 6.5°, with horizontal spacing centers at approximately −23°, −14.1°, −4.7°, +4.7°, +14.1°, and +23° relative to screen center. Vertically, the row was positioned 2.5° above the screen center.

In the CCMT, car study images measured 741 × 750 pixels and were embedded within frames subtending approximately 12.3° (width) and 5.7° of visual angle. During test trials, three cars were arranged in a vertically staggered column, with each subsequent car offset by ~4.6° of visual angle.

#### 3.2 Bayesian analyses: Robustness

Robustness was assessed by re-running all analyses across a range of prior widths (1–10). Corresponding BF robustness plots are provided in the supplementary materials (section 6), with varying prior widths (tested from 0 to 1.5, corresponding to the default range, and up to 10). We did not find any deviations from the primary results.

##### 3.3 Bayesian analyses: Robustness plots

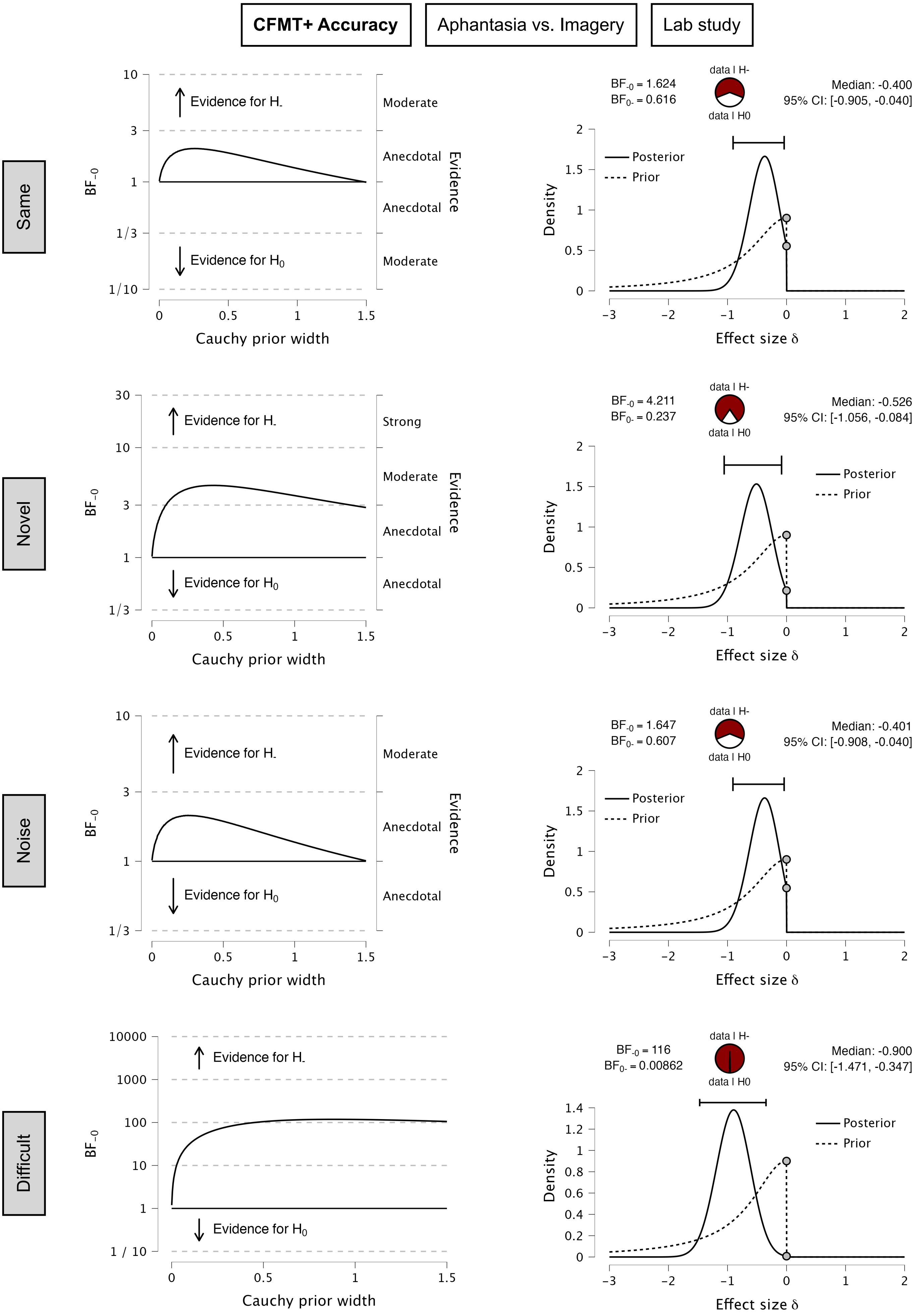


*Note*. Left panel: Robustness analysis of the Bayes Factor ($BF_{-0}$) as a function of the Cauchy prior width. Right panel: Prior (dashed line) and posterior (solid line) distributions for the effect size δ.

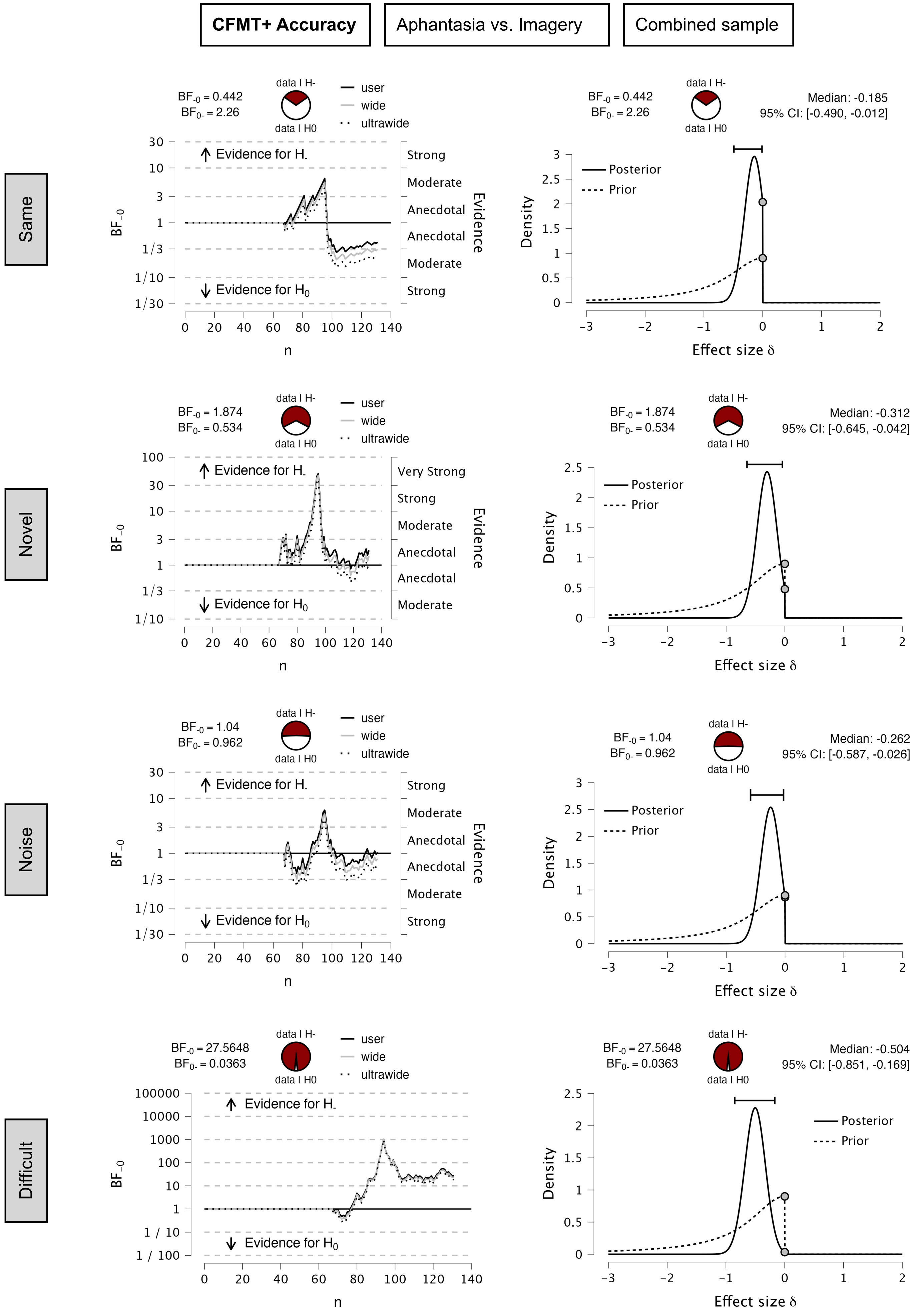


*Note*. Left panel: Robustness analysis of the Bayes Factor ($BF_{-0}$) as a function of the Cauchy prior width. Right panel: Prior (dashed line) and posterior (solid line) distributions for the effect size δ.

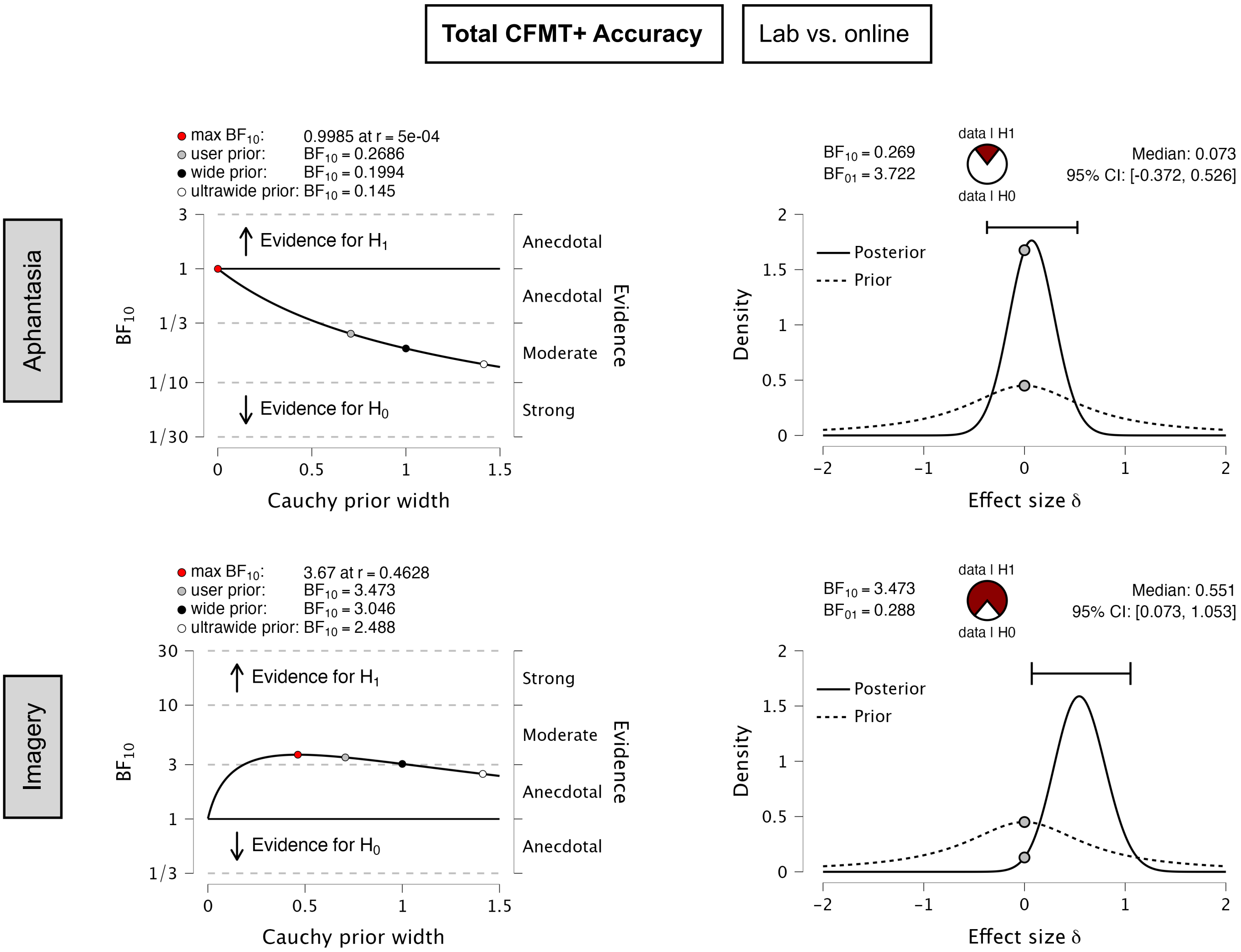


*Note*. Left panel: Robustness analysis of the Bayes Factor ($BF_{10}$) as a function of the Cauchy prior width. Right panel: Prior (dashed line) and posterior (solid line) distributions for the effect size δ.

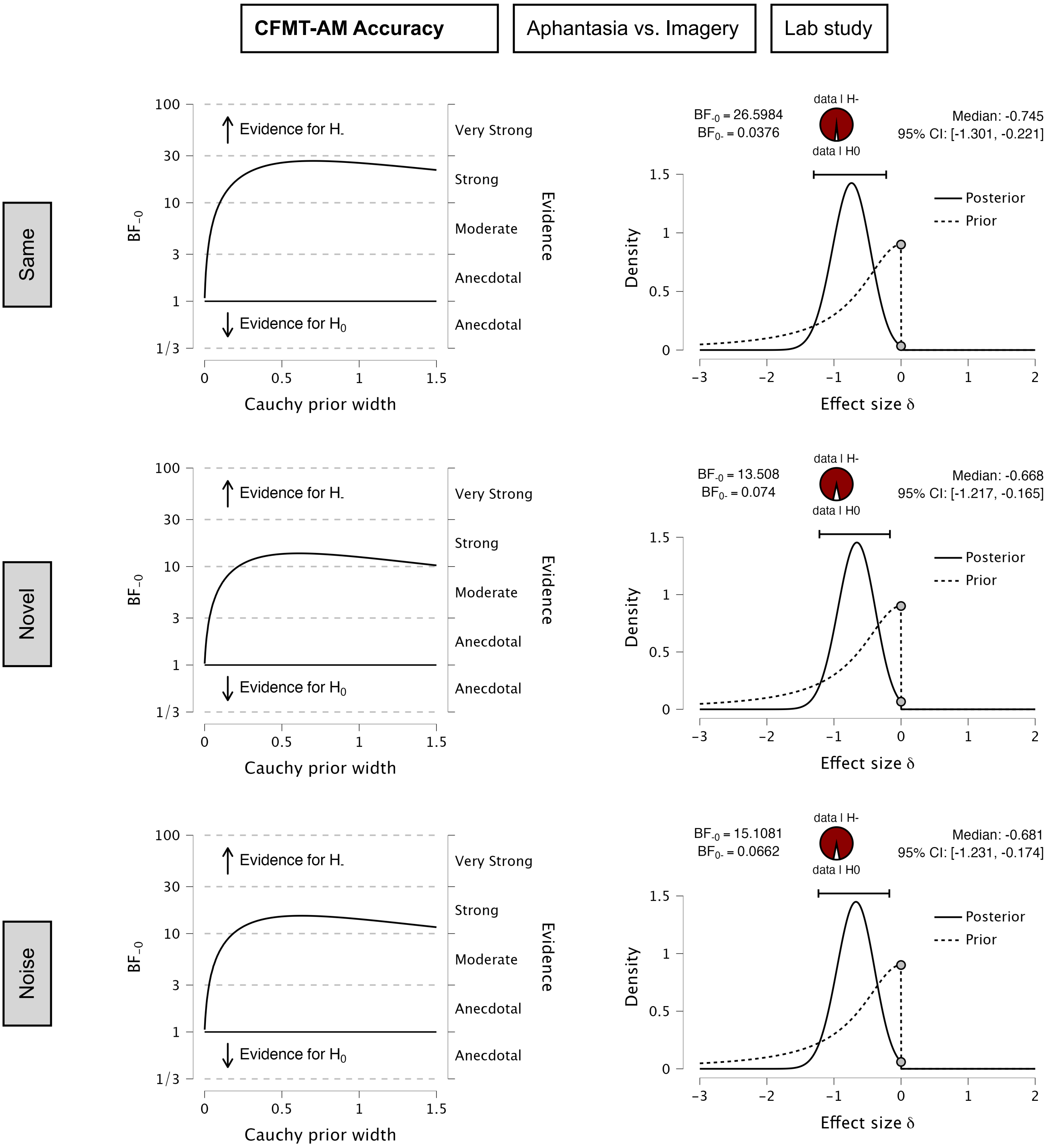


*Note*. Left panel: Robustness analysis of the Bayes Factor ($BF_{-0}$) as a function of the Cauchy prior width. Right panel: Prior (dashed line) and posterior (solid line) distributions for the effect size δ.

**RT slopes**

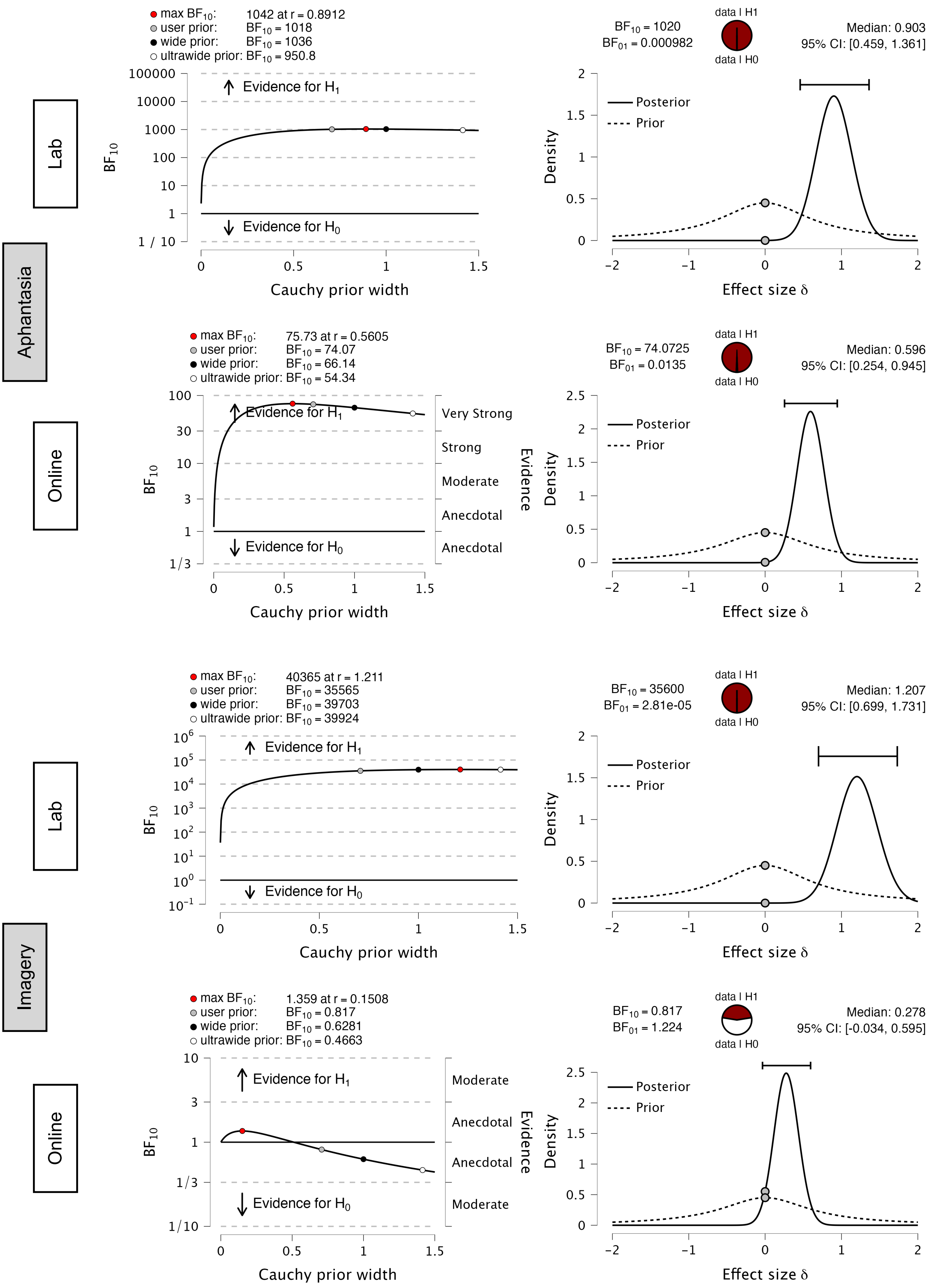


*Note*. Analysis of differences in RTs across levels of item difficulty per group and cohort. Left panel: Robustness analysis of the Bayes Factor ($BF_{10}$) as a function of the Cauchy prior width. Right panel: Prior (dashed line) and posterior (solid line) distributions for the effect size δ.

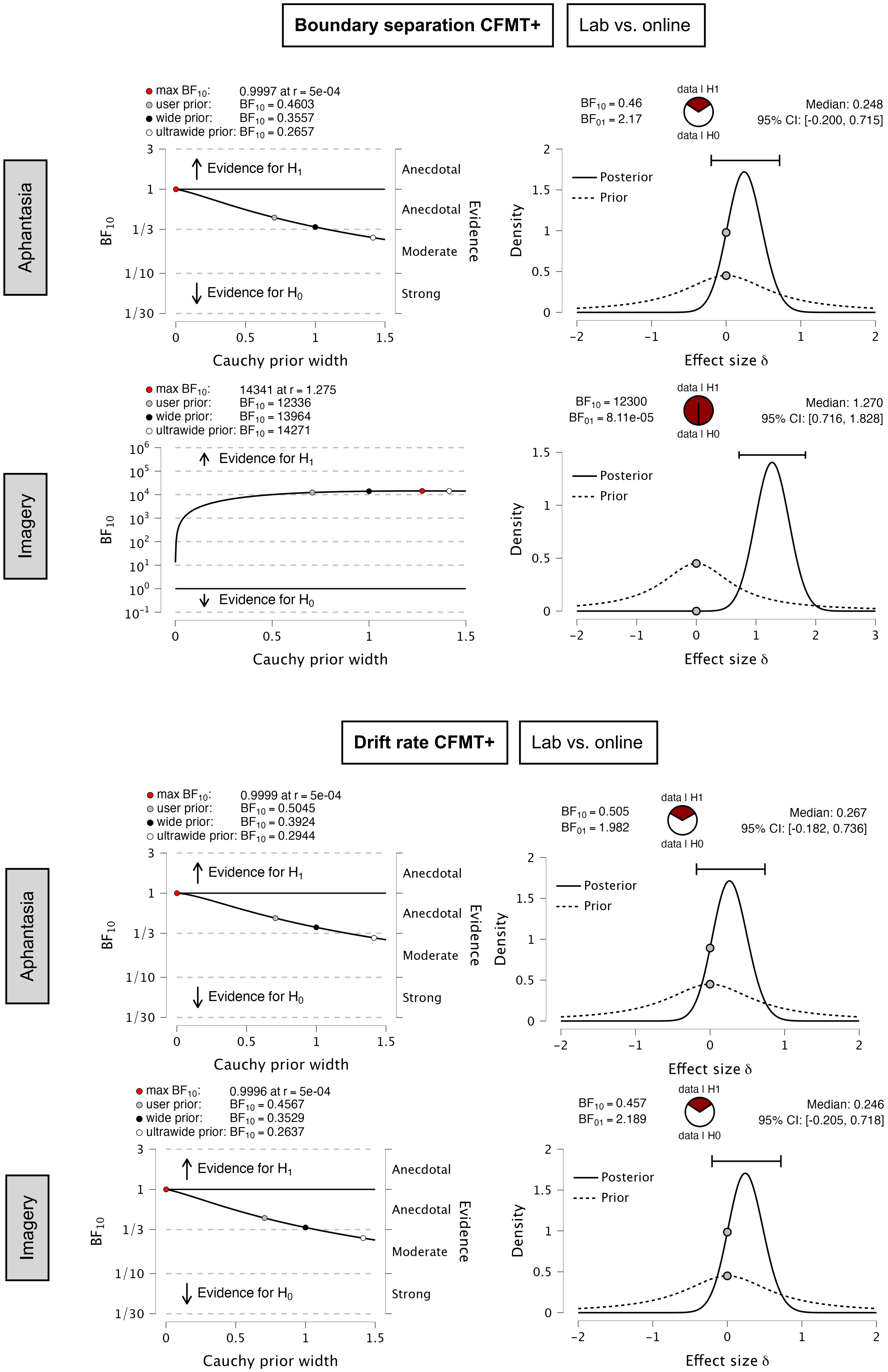


*Note*. Analysis of differences in RTs across levels of item difficulty per group and cohort. Left panel: Robustness analysis of the Bayes Factor ($BF_{10}$) as a function of the Cauchy prior width. Right panel: Prior (dashed line) and posterior (solid line) distributions for the effect size δ.

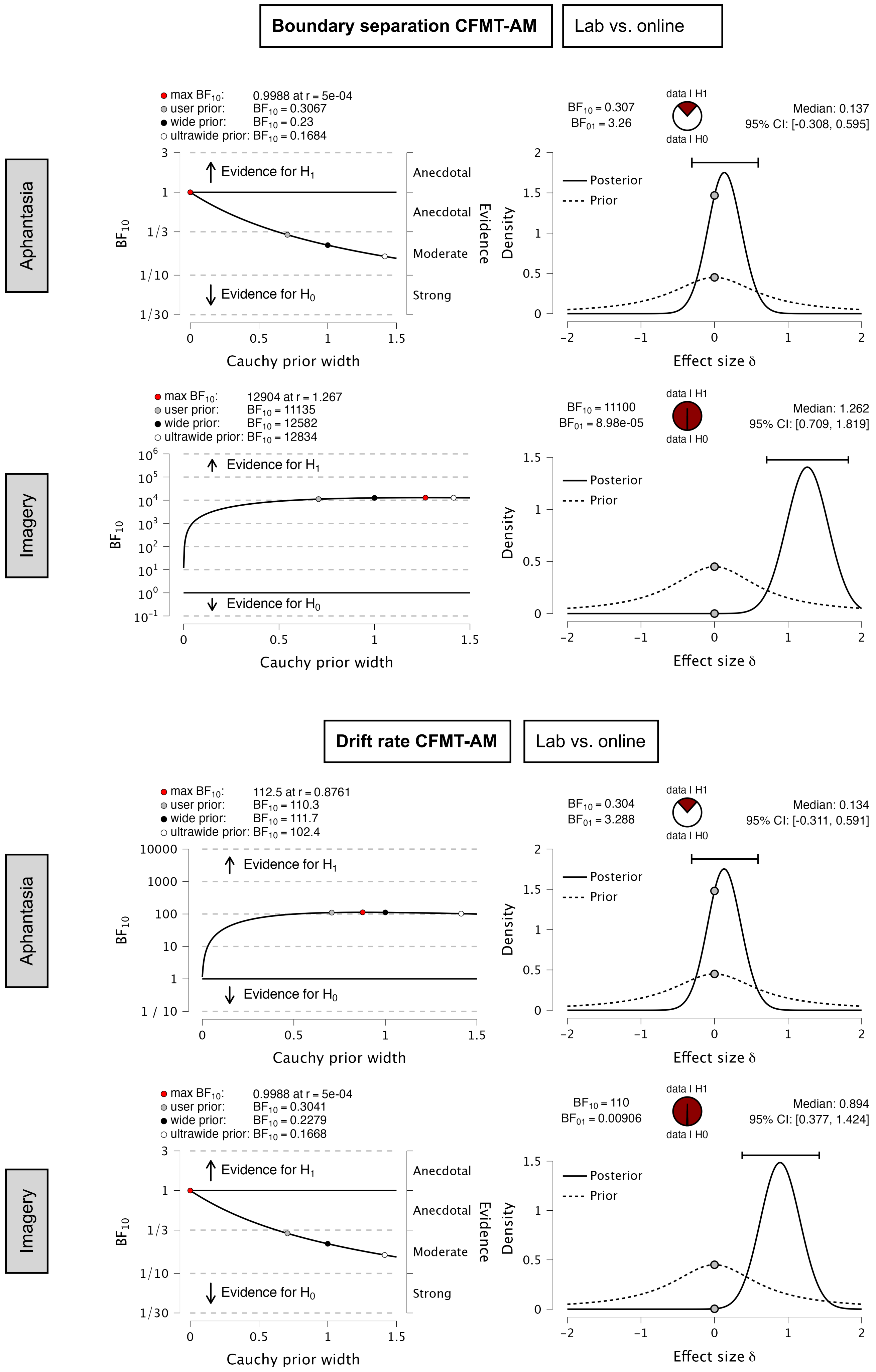


*Note*. Analysis of differences in RTs across levels of item difficulty per group and cohort. Left panel: Robustness analysis of the Bayes Factor ($BF_{10}$) as a function of the Cauchy prior width. Right panel: Prior (dashed line) and posterior (solid line) distributions for the effect size δ.

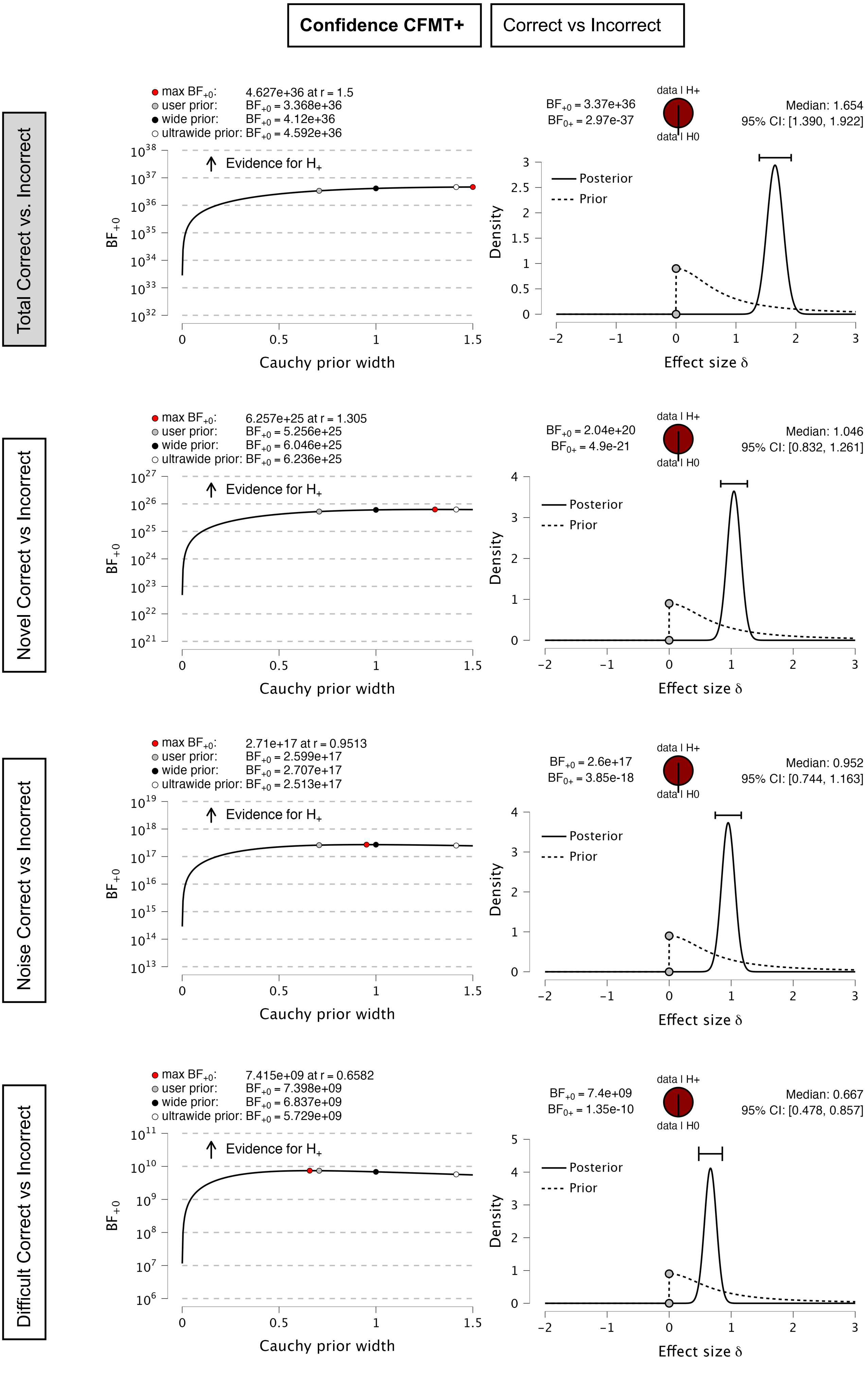


*Note*. Analysis of differences in RTs across levels of item difficulty per group and cohort. Left panel: Robustness analysis of the Bayes Factor ($BF_{10}$) as a function of the Cauchy prior width. Right panel: Prior (dashed line) and posterior (solid line) distributions for the effect size δ.

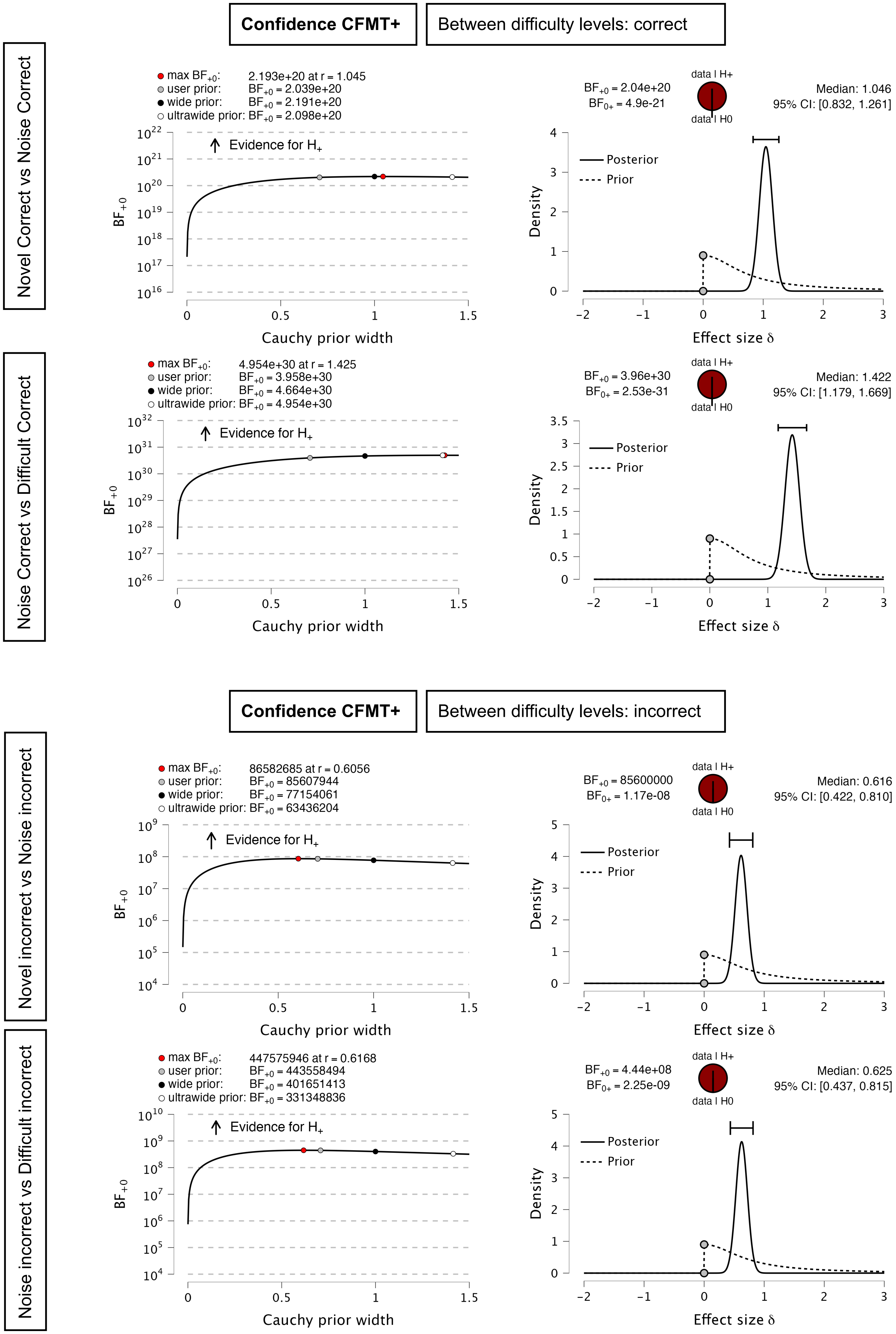


*Note*. Analysis of differences in RTs across levels of item difficulty per group and cohort. Left panel: Robustness analysis of the Bayes Factor ($BF_{10}$) as a function of the Cauchy prior width. Right panel: Prior (dashed line) and posterior (solid line) distributions for the effect size δ.

**Confidence CFMT+** | Between difficulty levels: correct - incorrect

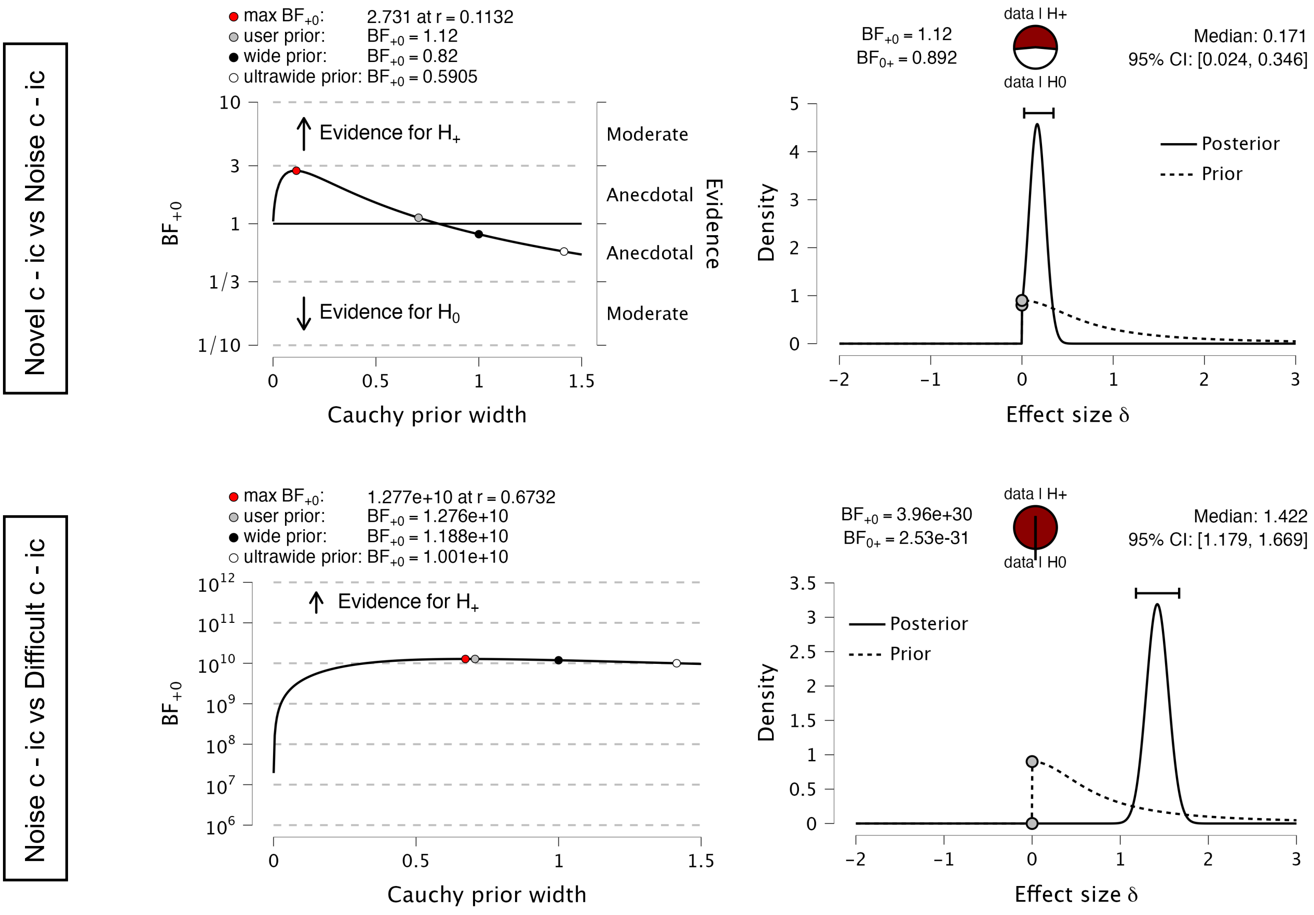


*Note*. Analysis of differences in RTs across levels of item difficulty per group and cohort. Left panel: Robustness analysis of the Bayes Factor ($BF_{10}$) as a function of the Cauchy prior width. Right panel: Prior (dashed line) and posterior (solid line) distributions for the effect size δ.
